# Detecting neural state transitions underlying event segmentation

**DOI:** 10.1101/2020.04.30.069989

**Authors:** Linda Geerligs, Marcel van Gerven, Umut Güçlü

**Affiliations:** Donders Institute for Brain, Cognition and Behaviour, Radboud University, Nijmegen, the Netherlands

**Keywords:** Event segmentation, fMRI, Hidden Markov Model, timescales, neural states, greedy search

## Abstract

Segmenting perceptual experience into meaningful events is a key cognitive process that helps us make sense of what is happening around us in the moment, as well as helping us recall past events. Nevertheless, little is known about the underlying neural mechanisms of the event segmentation process. Recent work has suggested that event segmentation can be linked to regional changes in neural activity patterns. Accurate methods for identifying such activity changes are important to allow further investigation of the neural basis of event segmentation and its link to the temporal processing hierarchy of the brain. In this study, we introduce a new set of elegant and simple methods to study these mechanisms. We introduce a method for identifying the boundaries between neural states in a brain area and a complementary one for identifying the number of neural states. Furthermore, we present the results of a comprehensive set of simulations and analyses of empirical fMRI data to provide guidelines for reliable estimation of neural states and show that our proposed methods outperform the current state-of-the-art in the literature. This methodological innovation will allow researchers to make headway in investigating the neural basis of event segmentation and information processing during naturalistic stimulation.

**Highlights:**

- Boundaries between meaningful events are related to neural state transitions.
- Neural states are temporarily stable regional brain activity patterns.
- We introduce novel methods for data-driven detection of neural state boundaries.
- These methods can identify the location and the number of neural state boundaries.
- Simulations and empirical data support the reliability and validity of our methods.

## Introduction

To understand the world around us as it unfolds over time, two processes are essential; information integration and segmentation. We integrate current sensory input with information from the past to make sense of speech or actions that unfold over time (Buonomano and Maass, 2009; Kiebel et al., 2008). We also segment information into distinct events when information from a previous timepoint is no longer relevant for what is occurring now (Kurby and Zacks, 2008; Newtson et al., 1977). Behavioral research has shown that segmenting information into meaningful events enables us to understand ongoing perceptual input (Zacks et al., 2001) and recall distinct events from our past (Flores et al., 2017; Sargent et al., 2013; Zacks et al., 2006). Although segmentation plays a fundamental role in the way we perceive and remember information in daily life, a lot remains unknown about the neural mechanisms that underlie these abilities. Here, we introduce a new method to investigate those mechanisms.

Recent innovations in data-driven analyses of functional magnetic resonance imaging (fMRI) voxel activity patterns during naturalistic stimulation have shown that event boundaries co-occur with shifts between stable patterns of brain activity (Baldassano et al., 2017). We will refer to these stable time periods as neural states, to distinguish them from the subjectively experienced events that have been described before (Zacks et al., 2007). By studying the duration of neural states, it is possible to see the temporal hierarchy of cortical information processing. High-level brain regions, such as the medial prefrontal cortex, show state durations that are comparable to those of experienced events. Lower-level brain areas such as visual cortex, show state durations at much shorter time-scales (Baldassano et al., 2017).

This observed temporal hierarchy is in line with findings from previous studies that have used different approaches to demonstrate a similar hierarchy for information integration in the cortex (Hasson et al., 2015; Honey et al., 2012; Lerner et al., 2011). These studies have shown that representations in lower level cortical areas quickly stabilize while representations in higher level brain regions are affected by information presented over thirty seconds before (Lerner et al., 2011). At first glance, this gradual build-up of representations may seem incompatible with the sudden shifts in brain activity patterns that are observed at neural states boundaries. However, recent work by Chien and Honey (2020) showed that both phenomena can be at play at the same time; while representations are constructed at varying rates across the cortex, context forgetting occurs at a similar rate. This suggests that neural state boundaries reflect context gating due to increases in local prediction error, which allows for removing irrelevant prior information at state boundaries (DuBrow et al., 2017; Ezzyat and Davachi, 2011). State transitions have also been shown to be coupled to a subsequent increase in activity in the hippocampus. This boundary-related hippocampal activity predicted reinstatements of brain activity during later recall, suggesting that state boundaries play an important role in memory encoding (Baldassano et al., 2017). In an electroencephalography (EEG) study, state transitions have been shown to be associated with a rapid reinstatement of the just-encoded movie state which may be associated with memory formation (Silva et al., 2019).

These results show that data-driven state detection methods are a valuable tool for investigating the neural mechanisms underlying event segmentation and memory formation as well as the timescales of cortical information processing. Although a lot of methods have been proposed to identify change points in functional connectivity patterns across large-scale brain networks (Cribben et al., 2012; Kundu et al., 2018; Xu and Lindquist, 2015), much less work has investigated state-transitions that are driven by changes in brain activity patterns within brain regions. The state detection method of Baldassano et al. (2017) is an exception. However, this method has not yet been systematically validated using simulated data. Here we investigate the accuracy and reliability of this hidden Markov model (HMM)-based state segmentation method and show a number of important limitations. To address these limitations, we introduce a much simpler greedy state boundary search (GSBS) method that outperforms the HMM-based method in terms of accuracy of state boundary detection and computational speed without making any assumptions about when state boundaries should occur. In addition, we introduce a novel t-distance metric for identifying the optimal number of states, which we also validate using simulations. We use empirical fMRI data to illustrate how the reliability of state boundary detection is improved with GSBS compared to the HMM-based approach. Empirical data also confirms that our new method for estimating the optimal number of states can recover the expected cortical temporal hierarchy. Finally, we use simulations and real data to explore how noise and data averaging might impact our ability to accurately identify state boundary transitions.

## Methods

### State segmentation methods

#### Greedy state boundary search method

The GSBS method we introduce here relies on a simple greedy search algorithm to identify the location of state boundaries. The state boundaries partition neural data into multiple time segments (clusters) that are each associated with a unique neural state (see figure 1A). The method searches for transitions between states, but is not designed to identify recurring states. State boundaries are identified in an iterative fashion, as described in more detail in the following.

**Figure 1:**
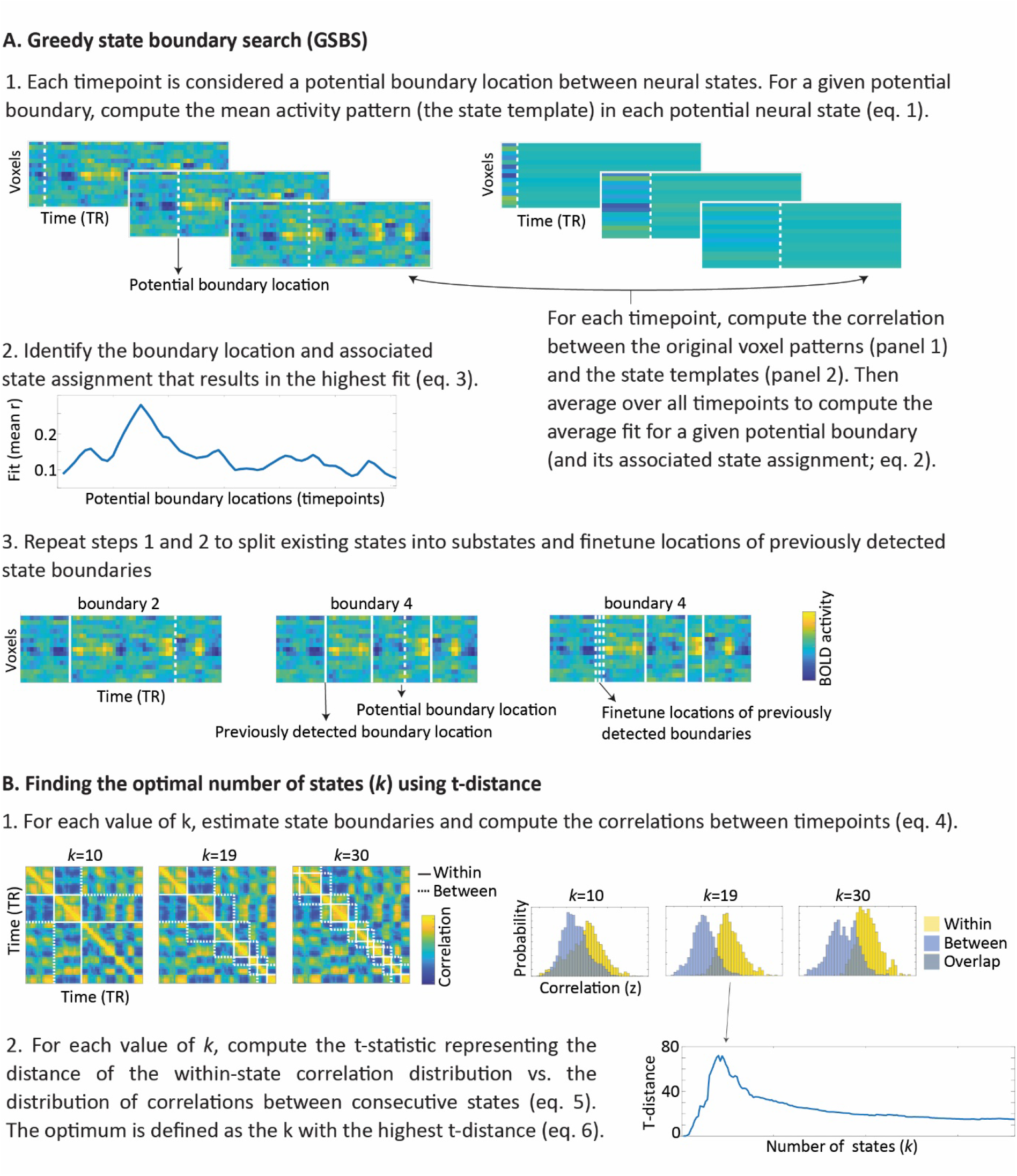
A) Illustration of the GSBS method that we use to identify state boundaries. B) Illustration of the t-distance metric that we use to determine the optimal number of states.

Let ***X*** be a *V* × *T* matrix of neural timeseries where *V* is the number of voxels and *T* is the number of timepoints. Let ***x***_*t*_ denote the *t*-th column of this matrix. Our approach boils down to a clustering algorithm where each timepoint t is assigned to exactly one state and members of a state are neighboring timepoints. Let ***S**_N_* = {*S*_1_,…, *S_N_*} denote the set of states identified after *N* iterations of the clustering algorithm, where the first timepoint within a given state is defined as the boundary location *b_n_* = min(*S_n_*). For each state *S_n_*, we use

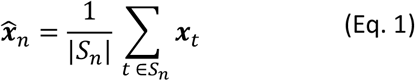

to denote the mean activity pattern over all timepoints with that state. GSBS aims to identify states that optimize the similarity between the mean activity patterns within states (the state template) and the original neural data at the corresponding timepoints. To this end, we define

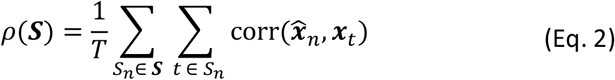

as the mean correlation between each timepoint and its state template.

Initially, all timepoints are in the same state and we initialize the set of state assignments as ***S***_1_ = {*S*_1_} with *S*_1_ = {1,…, *T*}. Next, in each iteration of the algorithm we split a state into two substates to obtain a more fine-grained partitioning of the timeseries. Let ***S**_N_*(*p*) denote the state assignment that is obtained by splitting a state *S_n_* ∈ ***S**_N_* with *p* ∈ *S_n_* into two potential substates 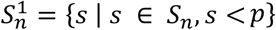 and 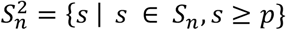, where *p* is the potential boundary location between these two potential substates.

At iteration N+1, we consider all timepoints as potential boundary locations and we identify the boundary location *p* and corresponding state assignment ***S**_N_*(*p*) that results in the maximal similarity between the state templates and the original neural data:

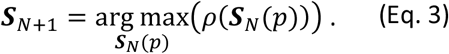

Since selecting a boundary can change the optimal location of the previous boundaries (Truong et al., 2020), we fine-tune all boundaries after a new boundary has been placed. One-by-one, the previously identified boundaries are shifted (± 1 TR) if this shift improves the fit, in the order in which the boundaries were initially identified. We constrained the potential shift to ± 1 TR because we observed that allowing a wider range did not further improve performance of the algorithm. The whole process of placing new boundaries and fine-tuning previous boundaries is repeated until a predefined number of states has been identified. When the number of states is unknown, our empirical observations suggest that it is sufficient to explore all possible number of states until it equals half the number of time points. This process leads to a set of boundary locations for each possible number of states, based on which the t-distance metric (described below) can be used to identify the optimal number of states.

#### Estimating the number of states with t-distance

While our GSBS method can identify state boundaries, it cannot determine the optimal number of states. Therefore, we designed a new metric to determine the optimal number of states *k*\*, which we call t-distance. T-distance is based on maximizing the similarity of timepoints in the same state, while minimizing the similarity of timepoints in consecutive states. The first step to compute the t-distance for a given number of states *k* is to compute the correlation between the neural activity patterns at each pair of timepoints *i, j* (see figure 1B, panel 1). That is,

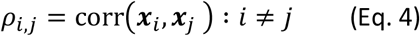

Pairs of timepoints that fall within the same state are considered to be part of the within-state correlation distribution and pairs of timepoints that are in consecutive states are part of the between-consecutive-state correlation distribution. The distance between the distribution of within-state correlation values *ρ_w_*(*k*) and the distribution of between-consecutive-state correlation values *ρ_b_*(*k*) is quantified using a t-statistic *t*_*ρ*_*w*_(*k*),*ρ_b_*(*k*)_ (see figure 1B, panel 2),

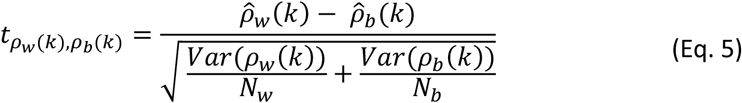

where *N_w_* is the number of within state correlation values and *N_b_* is the number of between-consecutive-state correlation values.

This t-distance is computed for each possible number of states and the optimal number of states is defined as the one with the highest t-distance. That is,

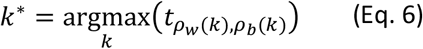

Note that by definition, the t-distance cannot be computed for k = 1 states. Code that implements our methods in Python is available in the StateSegmentation Python package (https://pypi.org/project/statesegmentation/). The code that was used to run the simulations and analyses shown in the paper can be found on Github (https://github.com/lgeerligs/State-segmentation-GSBS).

#### Baseline methods

To evaluate the performance of our GSBS method for detecting state boundaries and the t-distance metric for determining the optimal number of states, we compare it with existing methods.

The baseline we use for the GSBS method is the ‘event segmentation model’ created by Baldassano et al. (2017), which is a variant of a Hidden Markov Model (HMM). We used the Python implementation in the Brain Imaging Analysis Kit (version 0.10). This HMM-based state boundary detection method models the brain activity as a sequence of hidden (unobserved) states. Each state is characterized by a specific mean activity pattern across voxels. In contrast to regular HMMs, this variant is constrained such that there are no recurring states (i.e. the HMM is *null recurrent*). Therefore, the first timepoint of a brain activity timecourse is always in state 1 and the final timepoint is always is state *k*, where *k* is the total number of states. From one timepoint to the next, a brain region can either stay in the same state or jump to the next state. In this implementation, all states are fixed to have the same probability of staying in the same state versus jumping to the next state. The inputs to the HMM-based method consists of a set of z-scored voxel timecourses and a value for *k* (i.e., the number of states that needs to be estimated).

Because of known issues with the HMM for neural states with uneven lengths, a recent update of the method introduces an additional optimization step (Baldassano, 2020). This step tries to find neighbouring pairs of states with very similar patterns, indicating that they should be merged. It also looks for states that could be split in half into two very different-looking states. It examines if the fit can be improved by simultaneously merging two of the states and splitting one of the states, to keep the same number of states overall. We will refer to this implementation of the HMM as HMM split-merge (HMM-sm).

To evaluate the performance of our t-distance metric, we compare it to metrics that have been used before. The metric that was introduced by Baldassano et al. (2017) is based on subtracting the average correlation between all states from the average correlation within states. The optimal number of states is defined as the number of states with the largest difference of within-versus across-state correlations (we will refer to this as WAC). The two crucial differences between the t-distance and WAC metrics are 1) t-distance uses the t-statistic while WAC uses the average difference and 2) t-distance only considers the correlations between consecutive states, while WAC averages correlations between all states. Another metric that we used as a baseline is the log likelihood (LL) that measures the HMM model fit (i.e. the likelihood of the data given the model). The latter metric cannot be applied to state boundaries obtained with GSBS. Although the HMM is fit to z-scored data, we found that the LL performed better as a metric for the optimal number of states when the non z-scored data was used, therefore the LL estimates in the paper are all derived from non z-scored data.

#### Simulation design

In analyses with real data, it is not possible to know the ground truth of state boundary locations. Therefore, we used simulations to determine how accurate our method is in recovering the simulated state boundaries under different circumstances. These simulations were an extended version of the toy simulations performed by Baldassano et al. (2017). In particular, we constructed state-structured datasets with *V* = 50 voxels and *T* = 200 timepoints and a TR of 2.47 seconds. The number of timepoints and the TR were selected to (approximately) match our real data. The number of voxels was set at 50 to ensure that there were enough voxels to compute a reliable correlation coefficient.

The number of states was varied between *k* = 5, *k* = 15 and *k* = 30 (default *k* = 15). To create state timeseries, we started by dividing the timeseries into equally long states. Next, to introduce variability in the state durations, the deviation between each initial and final state boundary location was determined by sampling from a uniform distribution ranging from −*q* to *q*, where 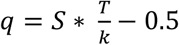, and *S* varied between 0.1, 1 and 2 (default *S* = 1). A mean pattern was drawn for each state from a standard normal distribution and the resulting voxel time courses were convolved with the canonical HRF from the SPM software package (default HRF peak delay = 6 s, peak dispersion = 1 s). To account for the initialization of the HRF, we initially created a longer timeseries (202 timepoints), these last two timepoints were always in the final state. After convolution, we removed the first 2 TRs from the time series. The simulated data for each timepoint and each voxel was the sum of the convolved state patterns plus a time course of equal length containing randomly distributed noise without autocorrelation with zero mean and standard deviation of SD=0.1. The noise SD was kept constant across simulated voxels. Each simulation was repeated 100 times, with different (randomly generated) state structures. An overview of the parameters used in each of the simulations can be found in table 1.

**Table 1:** Overview of the simulation parameters used to generate each of the figures

| Sim | Figure | Number of participants | Number of states | SD of state length | SD of random noise | Prob. of non-shared boundaries | HRF delay | HRF disp. |
| --- | --- | --- | --- | --- | --- | --- | --- | --- |
| A | 2 & S1 | 1 | 15/30 | 0.1/1/2 | 0.1 | 0 | 6 | 1 |
| B | 3 & S2 | 1 | 5/15/30 | 1 | 0.1 | 0 | 6 | 1 |
| C | 4 | 1 | 15 | 1 | 0.1 | 0 | 6 | 1 |
| D | 7 | 20 | 15 | 1 | 1/5/10 | 0 | 6 | 1 |
| E | 8 | 20 | 15 | 1 | 0.1 | 0.1/0.2/0.4 | 6 | 1 |
| F | S4 | 1 | 15 | 1 | 0.1 | 0 | 4/6/8 | 0.5/1/2 |

*Sim=simulation; prob=probability; disp= dispersion.*

To compare the compute time between the GSBS and the HMM-based methods, we simulated data for one participant in line with the details specified above. Then we measured the time that was needed to identify *k* neural states, where we would either loop through all possible states (assuming that *k* was unknown) or we only computed k states (assuming that *k* was known).

To investigate how the state boundary detection and the estimation of the optimal number of states is affected by noise in the data, individual differences, and averaging, we also simulated a group study. In these stimulations, we created datasets in which a group of 20 participants shared (some of) the states. In the first simulation, participants shared all states and state transitions and we varied the amount of random noise that was added to the data (between *SD* = 1, *SD* = 5 and *SD* = 10, in contrast to the *SD* = 0.1 specified above). The amount of noise was kept constant across all simulated participants. Then we investigated the effect of noise on state boundary detection and the estimation of the optimal number of states. We also investigated whether these effects of noise could be mitigated by averaging the data, or by using cross-validation, such that the state boundaries are defined in the training set and the optimal number of states are defined based on the test set. Data were averaged across all or half of the participants or cross validation was performed across 2 independent participant groups (folds) or using leave-one-out (LOO) cross validation. To investigate how averaging and cross–validation affect the results when not all states are shared between participants, we simulated data in which there is a specific probability that a given state transition that was present in the group, was not present in an individual, with *p* ∈ {0.1, 0.2, 0.4}. At the same time, we randomly added new state transitions to individuals, such that on average, across simulations, the number of states in each individual was the same as the number of states on the group-level. When a state transition that was present in the group disappeared in an individual, we modeled this by continuing the voxel pattern of the previous state. When we added individual-specific states, we generated a new unique mean activity pattern for each new state in each participant.

#### Measuring the similarity between state timeseries

To quantify the similarity between the two state timeseries (e.g. simulated and estimated states), we computed the Jaccard index; the number of timepoints that are in the same state divided by the total number of timepoints *T*. Let *k* denote the maximal number of states across both timeseries. We define a *k* × *k* matrix *M* such that *m_ij_* is the number of timepoints that are in state *i* in the simulated timeseries and in state *j* in the estimated timeseries. To be able to count the number of timepoints which are in the same state, state labels were first matched using the Hungarian method (Kuhn, 1955). The Hungarian method finds permutation matrices *L* and *R* such that the Jaccard index is given by:

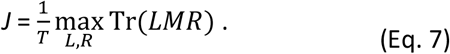

If the number of states in one timeseries is larger than in another, dummy rows or columns with zeroes are added to *M* to make it a square matrix. This implies that when the number of states are not equal across state timeseries, some state labels cannot be matched and therefore the Jaccard index can never be one.

Because the Jaccard index is affected by the number of states in both timeseries, we scaled it with respect to the expected similarity (i.e. the overlap that is expected by chance). This expected similarity was computed by generating 1000 random segmentations with the same number of states, in which the state boundary time points were uniformly sampled from the set of all time points (without replacement), and recomputing the Jaccard index (after matching the labels). The final similarity measure was scaled such that 0 indicates that the similarity is the same as the average expected similarity and 1 indicates perfect correspondence between the two state timecourses. Throughout the paper, we will refer to this measure as the adjusted accuracy. To further investigate the differences between simulated and estimated states, we also computed the distance between each estimated state boundary and the nearest simulated state boundary.

#### Dataset

We also used a real dataset to investigate performance of the different state boundary detection methods. In particular, we used 265 adults (131 female) who were aged 18–50 (mean age 36.3, SD = 8.6) from the healthy, population-derived cohort tested in Stage II of the Cam-CAN project (Shafto et al., 2014; Taylor et al., 2017). Participants had normal or corrected-to-normal vision and hearing, were native English speakers, and had no neurological disorders. Ethical approval for the study was obtained from the Cambridgeshire 2 (now East of England - Cambridge Central) Research Ethics Committee. Participants gave written informed consent.

Participants were scanned using fMRI while they watched a shortened version of a black and white television drama by Alfred Hitchcock called ‘Bang! You’re Dead’. In previous studies, a longer version of this movie has been shown to elicit robust brain activity, synchronized across younger participants (Hasson et al., 2009). Because of time constraints, the full 25-minute episode was condensed to 8 minutes with the narrative of the episode preserved (Shafto et al., 2014). Participants were instructed to watch, listen, and pay attention to the movie.

#### fMRI data acquisition & pre-processing

The details of the fMRI data acquisition are described in (Geerligs et al., 2018). In short, 193 volumes of movie data were acquired with a 32-channel head-coil, using a multi-echo, T2*-weighted echo-planar imaging (EPI) sequence. Each volume contained 32 axial slices (acquired in descending order), with slice thickness of 3.7 mm and interslice gap of 20% (repetition time (TR) = 2470 ms; five echoes [TE = 9.4 ms, 21.2 ms, 33 ms, 45 ms, 57 ms]; flip angle = 78 degrees; field-of-view = 192mm x 192 mm; voxel-size = 3 mm x 3 mm x 4.44 mm), the acquisition time was 8 minutes and 13 seconds. High-resolution (1 mm x 1 mm x 1 mm) T1 and T2-weighted images were also acquired.

The initial steps of data preprocessing were the same as in Geerligs et al. (2018) and are described there in detail. In short, the preprocessing steps included deobliquing of each echo time (TE), slice time correction and realignment of each TE to the first TE in the run, using AFNI (version AFNI_17.1.01; https://afni.nimh.nih.gov; Cox, 1996). Then multi-echo independent component analysis (ME-ICA) was used to denoise the data for each participant, facilitating the removal of non-BOLD components from the fMRI data, including effects of head motion (Kundu et al., 2012). Co-registration followed by DARTEL intersubject alignment was used to align participants to MNI space using SPM12 software (http://www.fil.ion.ucl.ac.uk/spm; Ashburner, 2007).

#### Hyperalignment

To optimally align voxels across participants in the movie dataset, we used whole-brain searchlight hyperalignment as implemented in the PyMVPA toolbox (Guntupalli et al., 2016; Hanke et al., 2009). Hyperalignment aligns participants to a common representational space based on their shared responses to the movie stimulus. Because the state boundary detection methods are applied to group averaged voxel-level data, good inter-subject alignment is essential. Hyperalignment uses Procrustes transformations to derive the optimal rotation parameters that minimize intersubject distances between responses to the same timepoints in the movie.

A common representational space was derived by applying hyperalignment iteratively. The first iteration started by hyperaligning one participant to a reference participant. This reference participant was chosen as the participant with the highest level of inter-participant synchrony across the whole cortex (i.e. strongest correlations between the participants’ timecourses and the average timecourses from the rest of the group, averaged across voxels). Next, a third participant was aligned to the mean response vectors of the first two participants. This hyperalignment and averaging alternation continued until all participants were aligned. In the second iteration, the transformation matrices were recalculated by hyperaligning each participant to the mean response vector from the first iteration. In a third iteration, the mean response vector was recomputed and this mean was defined as the common space. We then recalculated the transformation matrices for each participant to this common space. To align the whole cortex, hyperalignment was performed in overlapping searchlights with a radius of three voxels and a stepsize of two voxels between each of the searchlight centers. The individual searchlights were aggregated into a single transformation matrix by averaging overlapping searchlight transformations. These aggregated transformation matrices were used to project each participant’s movie fMRI data into the common representational space.

Note that hyperalignment and the analyses were performed on the same dataset. This approach might be an issue for classification analyses, due to the sharing of information between the training and test set. However, it is not a concern here because hyperalignment cannot influence the variable of interest, which is the temporal structure of the data.

#### Real data analyses

To investigate the performance of the different state segmentation methods on our data, we selected five regions of interest in the left hemisphere; V1 (MNI, x=-4, y=-90, z=-2), V5 (MNI, x=-44, y=-72, z=2, inferior temporal cortex (IT, MNI x=-50, y= −52, z=-8), angular gyrus (AG, MNI x=-42,y=-64,z=40) and medial prefrontal cortex (mPFC, MNI x=0, y=54, z=22). These peak coordinates were based on a search of these brain regions in the Neurosynth database (Yarkoni et al., 2011). Based on previous work, we expected to see a clear temporal hierarchy across these regions of interest, with the largest number of states in V1, which decreased in number as we move up cortical hierarchy to secondary visual processing areas (V5) and multi-modal association areas (AG & IT) toward the top of the cortical hierarchy in the mPFC (Baldassano et al., 2017; Hasson et al., 2015; Honey et al., 2012; Lerner et al., 2011).

Around each of these peak coordinates, we created spherical searchlights with different sizes (radius 6, 8, 10 or 12 mm, default = 8 mm). For the default of 8mm, each ROI contained approximately 80 voxels (range 72-82 voxels). We applied the different state segmentation methods in searchlights to the hyperaligned movie data. We observed that single-participant data was too noisy to reliably identify neural state boundaries. Therefore, we reduced the effects of noise by averaging the voxel timecourses across groups of participants with varying sizes; data were divided into either 2, 5, 10, 15, 20 or 265 distinct (groups of) participants, resulting in no averaging, or averages of, ~13, ~18, ~26, ~53 or ~128 participants. The default for the analyses was 15 groups of ~18 averaged participants, so that we had sufficient independent observations to compare the reliability across methods while reducing the noise effects as much as possible.

As in the simulations, we first aimed to establish the impact of the different state segmentation methods on the reliability of the state boundaries. Reliability was defined as the consistency of state boundaries, where the number of states was the same across the participant groups. The number of states was set to the optimal number of states averaged across all participant groups, as defined by GSBS and t-distance or defined apriori (k=20, 30 or 40). This was done to ensure that variability in the estimation of the number of states was not conflated with variability in the estimation of the location of state boundaries (which is shown separately). Reliability was estimated using the adjusted accuracy metric (as described in ‘Measuring the similarity between state timeseries’). The neural states in each group of participants can be described by a state labelled timecourse (timepoints numbered by state) and its corresponding state boundary timecourse (0 for no state transition, 1 for a state transition). The state labeled timecourse of one group of participants, with *k* states, was compared to a state labeled timecourse that was constructed based on the averaged state boundary timecourses across all other participant groups. This average state boundary timecourse across groups was converted to a state labelled timecourse by identifying the *k*-1 timepoints that were most often associated with a state transition and using these k-1 timepoints as state boundaries.

Besides investigating reliability, we also compared the different methods to estimate the optimal number of states in the empirical data. Specifically, we compared how the estimated optimal number of states derived from each these metrics aligns with the expected cortical hierarchy. To this end we combined the different boundary detection methods (GSBS/HMM) and fit metrics (t-distance/LL/WAC) of interest across all 15 groups. It has previously been suggested that LL should be used in combination with cross-validation (Chang et al., 2020). Therefore, we additionally investigated LL performance with cross validation, such that the averaged data of ~248 participants (14 groups) was used to fit the HMM and the data from the left out group was used to compute the LL.

As a third metric of interest, we compared the neural states transitions to the timing of actual event transitions as estimated by humans, we used subjective annotations on the occurrence of event boundaries in the Cam-CAN movie dataset that were collected by Ben-Yakov and Henson (2018). These annotations were based on sixteen observers who watched the movie outside the scanner and indicated with a keypress when they felt “one event (meaningful unit) ended and another began”. Only boundaries identified by a minimum of five observers were included, resulting in a total of 19 boundaries separated by 6.5–93.7 s. To create an event labeled timecourse, that could be compared to the neural state timecourses, boundaries were shifted with 5 seconds to account for the HRF delay and then TRs containing an event boundary were identified.

Statistical comparisons between the different state segmentation methods for the different metrics of interest were performed with the Wilcoxon signed rank test.

In a set of additional analyses, we investigated several factors that may affect the performance of the GSBS and t-distance metrics. First, averaging data across participants may reduce noise in the data, so we investigated the effects of data averaging on the reliability and the estimated optimal number of states. Second, because GSBS requires sufficient voxels in an ROI to be able to compute a reliable estimate of the correlation coefficient, we investigated the effect of the size of the searchlight sphere on the reliability and the estimated optimal number of states. Third, we investigated how hyperalignment and excluding voxels with low inter-subject synchrony (as in Baldassano et al., 2017) affected the reliability of neural states. Inter-subject synchrony was measured as the Pearson correlation between the time series of each participant and the average timeseries of all other participants. This estimate was averaged across all participants to obtain a group-average measure of inter-subject synchrony.

## Results

In the first simulation (simulation A in table 1), we evaluated the performance of the greedy state boundary search (GSBS) in detecting state boundaries in simulated data. As a baseline for our evaluation, we contrasted it with the HMM-based event-segmentation model (Baldassano et al., 2017). We used both the original implementation of the HMM-based method as well as a more recent version, which includes an additional splitting and merging step to optimize the solutions for uneven state lengths (HMM-sm). In this first simulation, we assume that the number of states is known (*k* = 15). To quantify the method’s performance, we computed the accuracy of state labels after an adjustment for the amount of overlap that would be expected by chance (adjusted accuracy; see methods). We found that when the simulated state boundaries are equally spaced (e.g. the standard deviation (SD) of state lengths is low), all three methods do a good job of recovering the states accurately (median adjusted accuracy = 1). However, when the states have variable lengths, we see that performance drops for the HMM-based methods but not for GSBS (see figure 2A). This performance drop is much more substantial for regular HMM than HMM-sm. Figure 2B shows the distribution of the distances between the simulated and estimated state boundaries. While the maximal distance for GSBS is 1 TR, the HMM-based methods show a range of distances. These distances are not uniformly distributed but show a peak around the middle (10-12 TRs distance), suggesting that the method has a tendency to recover states with similar lengths (the expected distance for equally spaced states is 13 TRs). These results were very similar for a different number of states (k=30, see supplementary figure 1). Together, these results show that the GSBS method performs better in recovering neural states, especially when states can vary in length.

**Figure 2:**
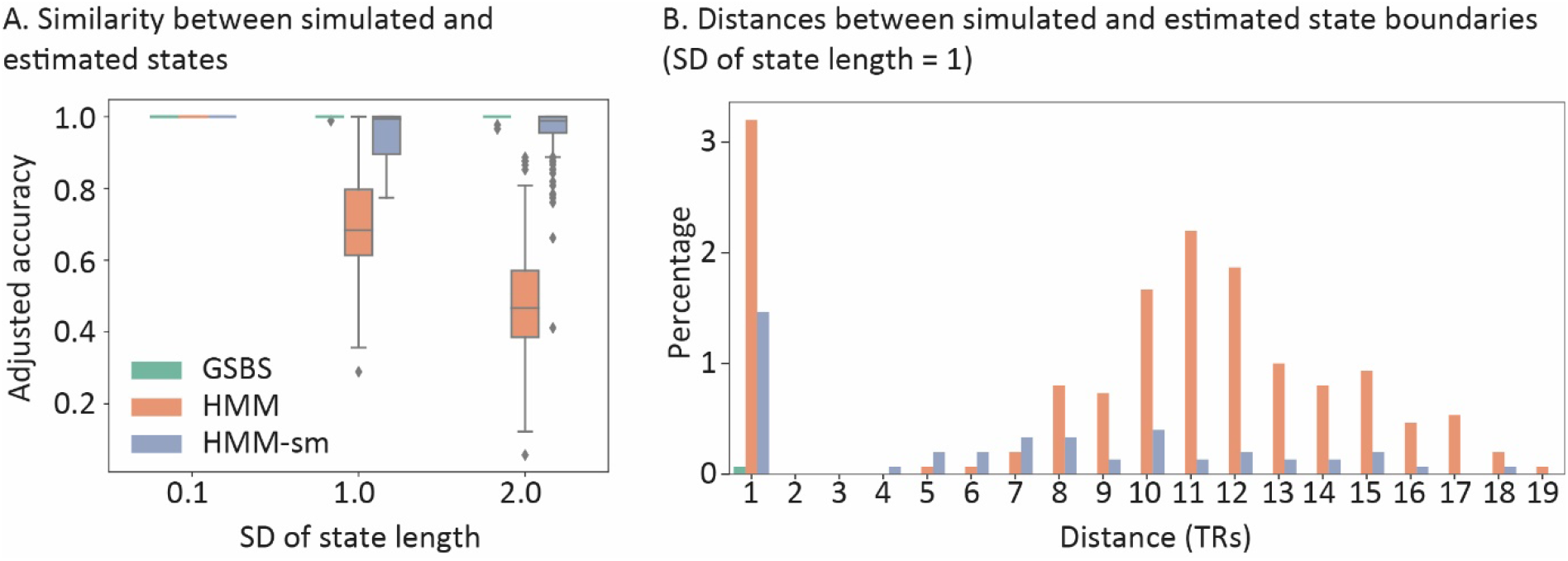
Results of simulation A. In this simulation the number of states is assumed to be known (k = 15). A) Similarity between simulated and estimated states for the HMM-based and GSBS methods. B) Distance between simulated and estimated state boundaries. The height of the bars indicates the percentage of the total number of estimated state boundaries for which the given distance was observed. Boundaries with a distance of zero (indicating perfect overlap between the simulated and estimates boundary) are not shown.

Simulation B was aimed at comparing the metrics used to estimate the optimal number of states. Specifically, we compared the t-distance metric (figure 1B), the WAC metric (introduced by Baldassano et al., 2017) and log likelihood (LL). T-distance uses the t-statistic to optimally separate the distributions of correlations within states and correlations between consecutive states. In contrast, WAC relies on optimizing the differences of within and between state correlations. WAC and t-distance can both be combined with the HMM-based method and GSBS. LL is a measure of overall model fit in the HMM and can therefore not be computed for GSBS. In line with previous recommendations, we used HMM (but not HMM-sm) to perform a search for the optimal number of states (Chang et al., 2020).

Figure 3A shows that when t-distance was combined with GSBS, the simulated number of states could be recovered quite accurately. There was a slight underestimation for *k* = 30, which may be because the method struggles to detect states with a duration of 1 TR. When t-distance was combined with HMM, the number of states was more strongly underestimated across all simulation parameters. When HMM was used in combination with LL, the number of states was underestimated for the simulation with 5 and 15 states, but correctly estimated for *k* = 30. WAC performed well for *k* = 5, but overestimated the number of states for *k* = 15 and *k* = 30, both for HMM and GSBS. We also investigated the adjusted accuracy of the state identification for this simulation with an unknown number of states. For all values of *k* we found that the states were recovered most accurately by GSBS combined with t-distance. Looking specifically at the HMM-based methods, we found best performance in combination with WAC and t-distance for k=5 and k=15, while for k=30, t-distance and LL gave the best performance. These results suggest that also for HMM-based methods, t-distance is the best metric to determine the optimal number of states.

**Figure 3.**
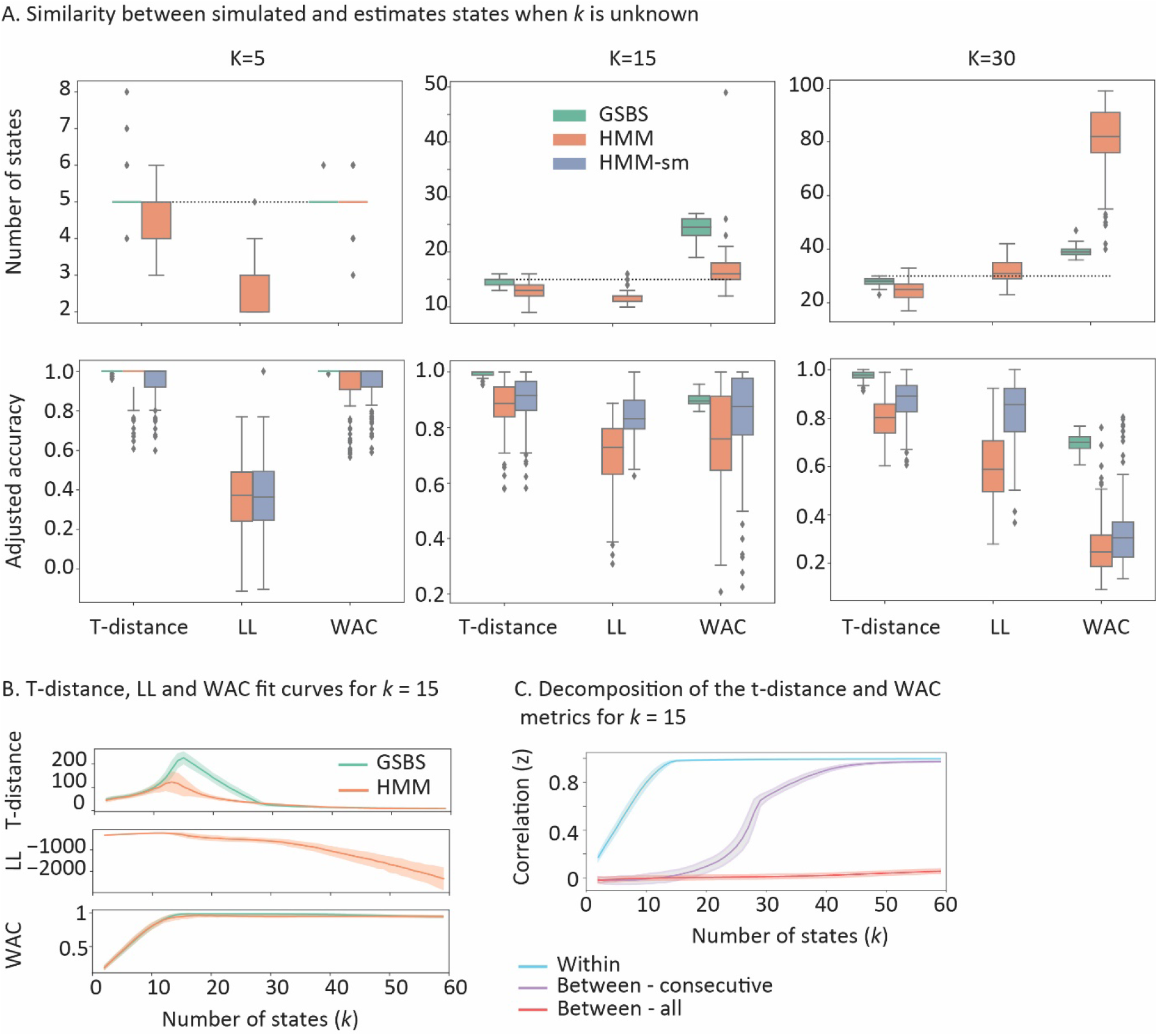
Results of simulation B. Comparison between the t-distance, LL and WAC metrics for estimating the optimal number of state boundaries in combination with HMM and GSBS. A) Comparison of the estimated and simulated number of states. The adjusted accuracy of each metric is computed for the estimated number of states shown in the top panel. In line with previous recommendations, we only used HMM (and not HMM-sm) to perform a search for the optimal number of states (Chang et al., 2020). For HMM-sm, the optimal number of states was set to the optimum estimated by HMM. B) Estimates of t-distance, LL and WAC for different number of states, in simulation where the real number of states (k) = 15. C) Comparison of the underlying components for the WAC and t-distance metrics for k = 15. WAC is computed by comparing the within state and the between – all correlation values. T-distance is based on the distribution of within state and between-consecutive state correlation values (see figure 1B). In panels B and C the shaded area indicates the standard deviation across simulation repetitions.

To examine the behaviour of the t-distance, WAC and LL metrics in more detail, figure 3B shows the plots of how the metrics vary across the number of estimated states (for simulation k=15), while figure 3C shows the elements that underlie the computation of the t-distance and WAC metrics. Comparing figures 3B and 3C shows that WAC is largely driven by within-state similarity, which increases slightly as more states are added even after the simulated number of states is exceeded, potentially due to autocorrelation caused by the HRF. Indeed, simulations without HRF convolution did not show this behavior. The between-state similarity hovers around zero regardless of the number of states that is estimated and does not have a lot of impact on WAC (in a simulation with 200 TRs). The overestimation of the number of states for the WAC method occurs even when four TRs around the diagonal are not taken into account (as in Baldassano et al., 2017, see supplementary figure 2). For t-distance, the same within-state similarity is used as for WAC. However, the ongoing increase in within-state similarity as more states are added is offset by the increase in similarity between consecutive states. This allows the method to identify the optimal number of states accurately. Both the within-state similarity and the similarity between consecutive states increase as the autocorrelation of the data increases, which may be why t-distance is less affected by HRF induced autocorrelation than WAC. LL increases initially and then declines steeply, even before the estimated number of states matched the simulated number of states. Together, these results show that the t-distance metric performs better in recovering the true number of states than the WAC and LL metrics, for both HMM and GSBS.

In simulation C, we compared the computational time that is required to run the GSBS and HMM-based methods for different numbers of states. When the number of states that should be estimated is known, the GSBS method results in a minimal improvement in computational speed compared to the regular HMM implementation but a 2-fold speedup compared to HMM-sm (see figure 4A). The differences between methods become more apparent when the number of states is not known and we need to go through all possible number of states to estimate the optimum. The HMM-based methods identify a new set of states for each value of *k* (the number of states). Therefore, the analyses need to be repeated for each value of *k*, which makes it computationally demanding. In contrast, the GSBS method performs an iterative search, which means that all but one of the boundaries that are detected for *k*=9 are based on the boundaries that are detected for *k* = 10 (although the location of previously detected boundaries is fine-tuned). These differences result in an up to 80-fold increase in computational speed for the GSBS method, compared to the basic HMM implementation and a 170-fold increase compared to the HMM-sm method (figure 4B). The results shown in figure 4 are for one simulated dataset with 200 timepoints and 50 voxels. When we are interested in investigating multiple searchlights across many participants and brain regions, the computational demands of the HMM-based methods quickly become prohibitive.

**Figure 4.**
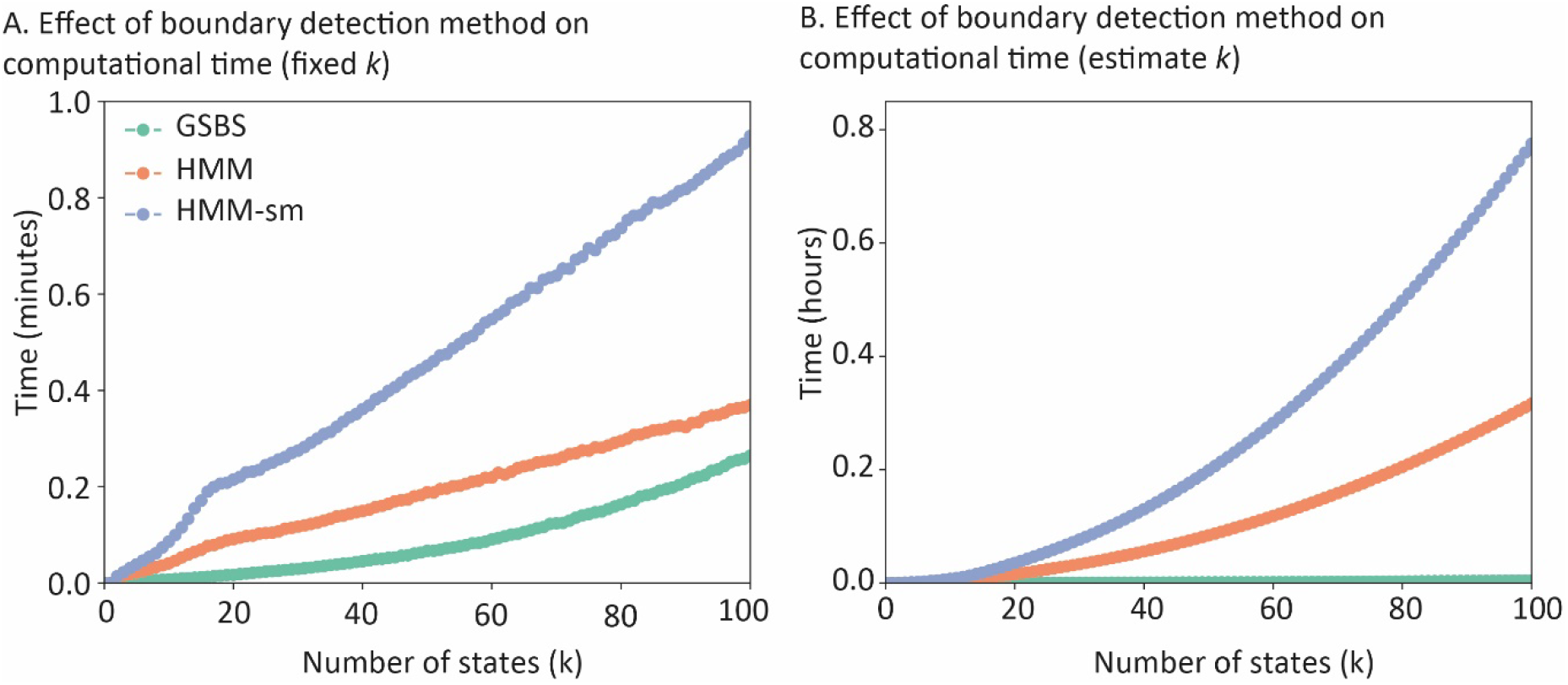
Results of simulation C. Comparison between the computational time required to run the GSBS and HMM-based boundary detection methods. A) The computational time when we assume that the number of states is known and is fixed at k. B) The computational time when we perform a search through all possible numbers of states, ranging from 2 to k. These results were obtained using a 3.0 GHz, 32 core CPU.

### Empirical data

Next, we compared the reliability of the state boundary detection for the GSBS and HMM-based methods for real fMRI data that was recorded while participants were watching a short movie. To be able to investigate the similarity of the results across participant groups as an estimate of reliability, participants were randomly divided into 15 groups of 17-18 participants. Within each group, the voxel timecourses were averaged within each ROI (the impact of averaging data will be explored in the next section). We investigated the reliability of the methods by computing the adjusted accuracy between the state timecourses in each group of participants and an average state timecourse derived from all other participant groups (similar to inter-subject synchronization). In addition, we estimated the ability of the t-distance, LL and WAC metrics to identify a plausible number of state boundaries. We also investigated LL performance with cross validation (in line with Chang et al., 2020). Based on previous work, we expected to see a clear temporal hierarchy across our five regions of interest (Baldassano et al., 2017; Hasson et al., 2015; Honey et al., 2012; Lerner et al., 2011). In particular, we expected the largest number of states in V1, which decreased in number as we move up the visual processing hierarchy (V5) and decreased further in multi-modal association areas, such as the angular gyrus and inferior temporal cortex (IT) and medial prefrontal cortex (mPFC).

Figure 5A shows that the GSBS method resulted in more reliable state boundaries across the 15 participant groups than the HMM and HMM-sm methods. There was one exception in area V5, where the HMM (but not HMM-sm) showed more reliable boundaries than GSBS. To derive the reliability estimates in figure 5A, we used the optimal number of states for each ROI, as determined by the t-distance combined with GSBS and shown in figure 6A. When we estimated different numbers of states (k=20, 30 or 40), we found that the reliability of the GSBS method was significantly higher than the reliability of one or both of the HMM-based methods for 10 out of 15 comparisons, while the HMM-based methods were more reliable in 2 comparisons (see supplementary figure 3). These results are in line with the simulation results, suggesting that the GSBS method outperforms the HMM-based methods in terms of reliably estimating state boundaries.

**Figure 5.**
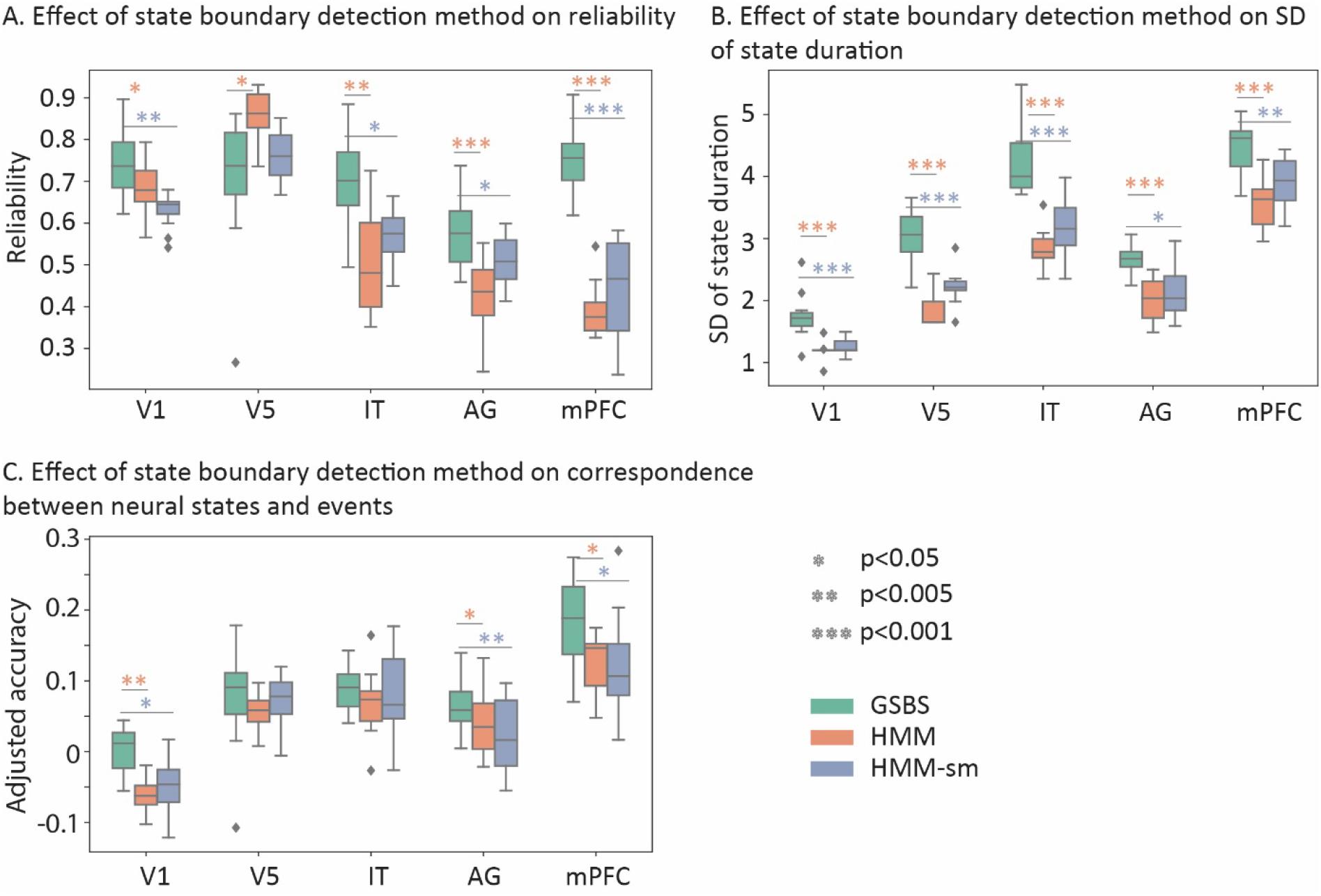
Comparison between the GSBS and HMM-based methods in real data. A) Reliability of state boundaries. B) SD of state boundary lengths. C) Similarity between neural states and events. The results in panels A-C were computed for the optimal number of states. This optimal number of states was determined by running GSBS for each possible number of states (k) and subsequently determining the optimal number of states using t-distance (results shown in figure 6A).

**Figure 6.**
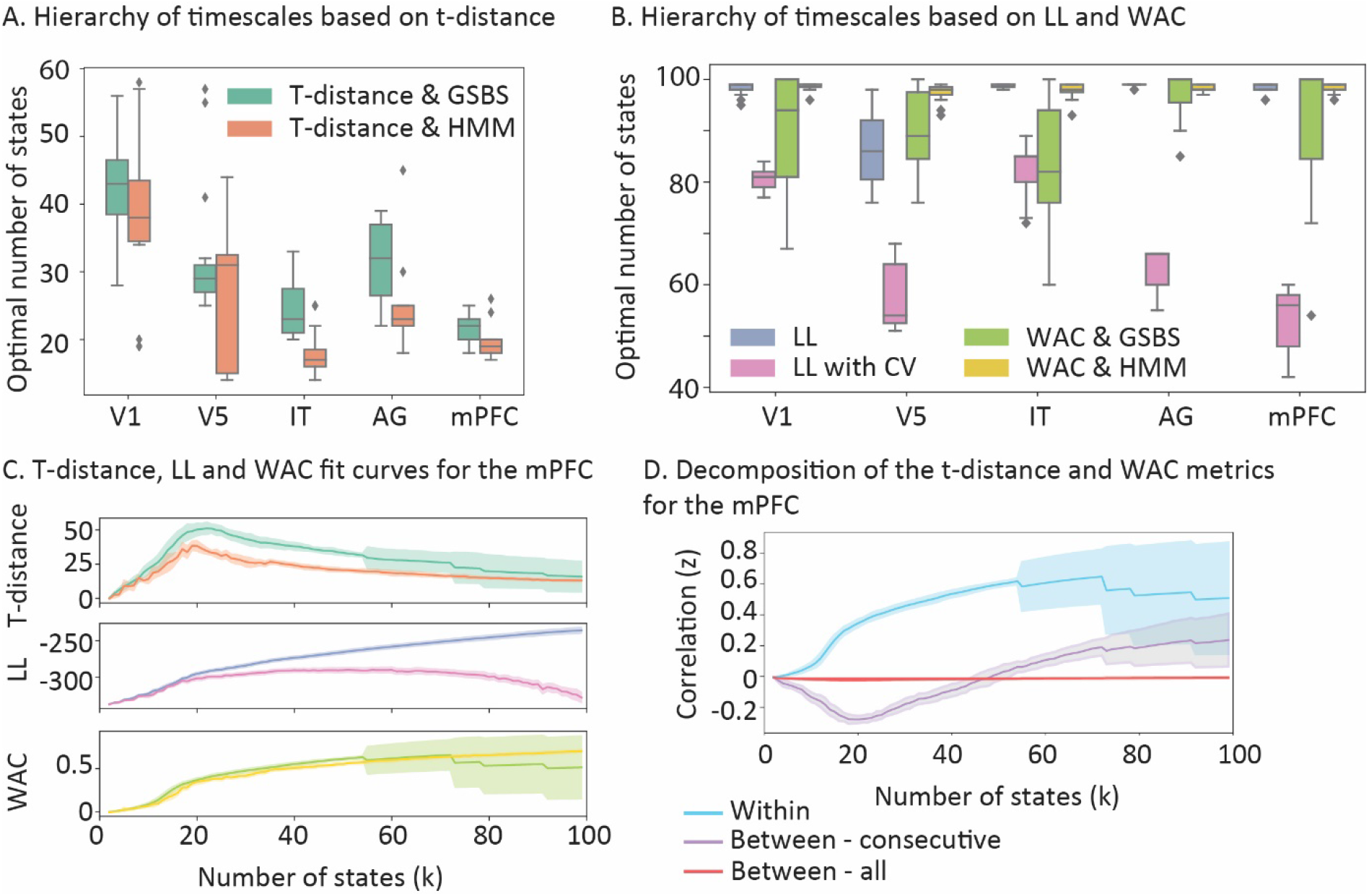
Comparison between the t-distance, WAC and LL metrics for real data. A) Optimal number of states for t-distance, combined with GSBS or HMM. B) Optimal number of states for WAC combined with GSBS or HMM or LL combined with HMM, with or without cross-validation. C) Estimates of t-distance, LL and WAC for different numbers of states in the mPFC. D) Comparison of the underlying components for the WAC and t-distance metrics in the mPFC. WAC is computed by comparing the within state and the between – all correlation values. T-distance is based on the distribution of within state and between-consecutive state correlation values (see figure 1B). In panels C and D the shaded area indicates the standard deviation across the 15 groups of participants.”

Simulation results suggested that the poorer performance of the HMM-based method is because it tends to estimate states that have a similar duration. If that is true, we would expect that states are more similar in length for the HMM-based than the GSBS method. Indeed, when we investigate the standard deviation of state lengths, we found that this was significantly higher for GSBS than the HMM-based methods for each of the ROIs (figure 5B), suggesting that the HMM-based method is biased toward finding states with similar durations. This deficit is not fully tackled by the split and merge step in the HMM-sm implementation.

We also investigated the similarity between the neural states and events derived from behavioral data. There, we found a significantly higher overlap between neural states and events for GSBS than the HMM-based methods in the angular gyrus and the mPFC (figure 5C). The mPFC was also the region that showed the highest overlap between events and neural states.

Figure 6A shows the estimates of the number of state boundaries, where the state boundaries were based on GSBS or HMM and the number of states were determined using t-distance. The estimated optimal number of states aligns well with the expected cortical hierarchy of timescales. In contrast, when we estimated the number of states using the WAC method (see figure 6B), we found an optimum of 80 states or more for each ROI (100 was the upper limit of the number of states we estimated). A similar pattern was observed for LL without cross validation. LL with cross validation did show results that aligned somewhat with the expected temporal hierarchy, although the number of states in each ROI was much higher than for t-distance. Specifically, the median number of states ranged from 54 to 85 for LL with cross validation combined with HMM, while it ranged from 22 to 43 for t-distance combined with GSBS and from 17 to 38 for t-distance combined with HMM.

Figures 6C and 6D illustrate why this happens. For WAC, the fit is only driven by the within-state similarity, while the between-state similarity stays around zero. Because the within-state similarity keeps increasing as more states are added, the optimal number of states is overestimated. For t-distance, the increase in within-state similarity is countered by the increase in the similarity between consecutive states, as the number of states increase. For LL, the fit keeps increasing as the number of states increase, which may be due to auto-correlated noise in the data. LL performs better when combined with cross validation, however in that case it still does not show a clear peak for the optimal number of states. These results are in line with the results for the simulated data, showing that t-distance provides a more accurate estimate of the optimal number of states than WAC and LL.

### Simulations – Assumptions, averaging and noise

Now that we have established that the GSBS method combined with the t-distance metric are the best tools to estimate state boundaries in fMRI data, our next step is to investigate the role of potential confounds on the estimation of the location and the number of state boundaries. First, we investigate the role of noise, which we simulate here as BOLD responses generated by brain activity without a particular state structure and we investigate how noise affects the estimation of the number of state boundaries. In simulation D, we simulate data from 20 participants who each move through the same 15 states and we add varying levels of noise to each participant’s data. When we identified state boundaries in each participant separately, we found that as the noise increases, the number of states was initially underestimated. However, as it increases further, the number of states was strongly overestimated (see light green bar in figure 7A). We also found that an increase in noise leads to a steady decline in the adjusted accuracy of the estimated state timecourses, when we assume that the number of states is known (see ‘no avg or CV’ in figure 7B).

**Figure 7.**
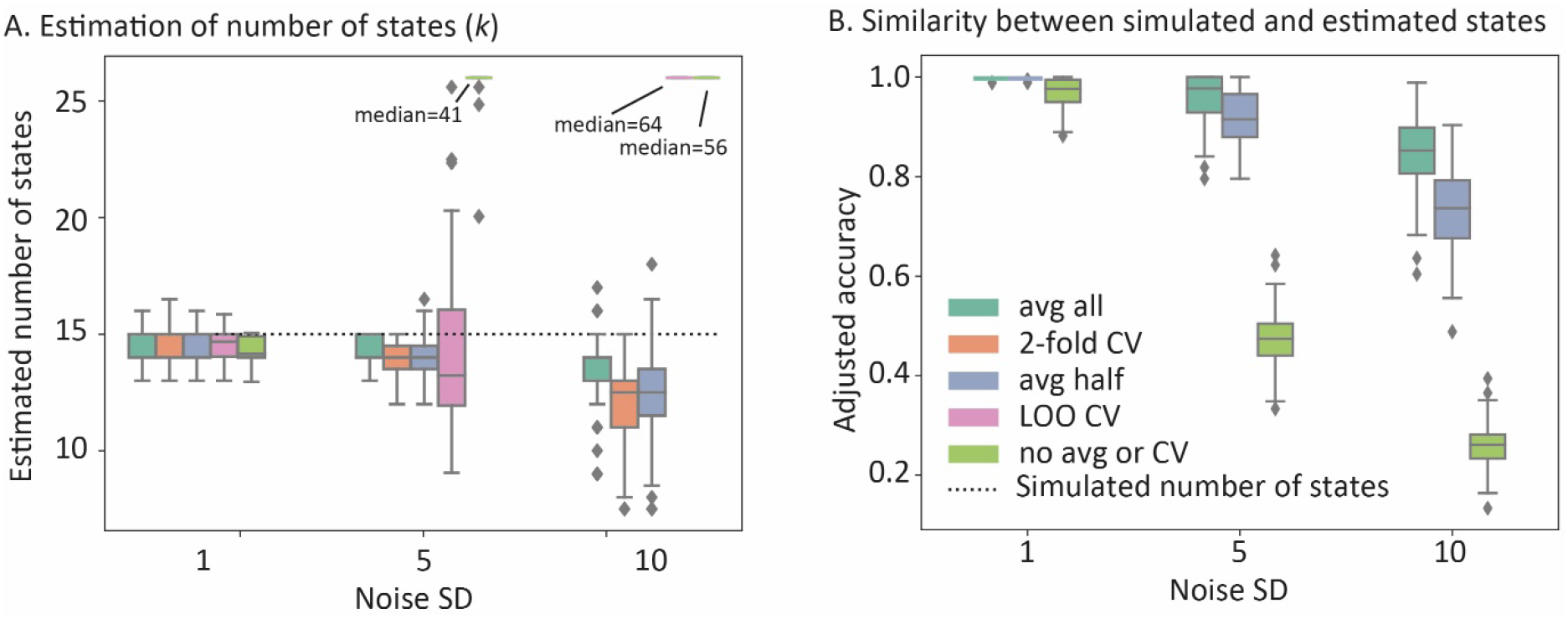
Results of simulation D. Comparing the effects of noise and data-averaging/cross validation on A) estimating the number of states B) the similarity between simulated and estimated state timecourses when we assume that the number of states (k=15) is known. avg all = averaging the data across all simulated participants; 2-fold CV = cross validation across 2 folds (independent groups). avg half = averaging the data in two independent groups; LOO CV = cross validation using the leave-one-out method; no avg or CV = using the data of simulated participants without averaging.

Next, we investigated two methods for reducing the impact of noise on the estimate of the number of states. One is to use cross-validation (as in Baldassano et al, 2017), in which the state boundaries are detected in one set of participants and the fit to determine the optimal number of states is derived from another set of participants. This reduces overfitting because the estimate of the number of states needs to be based on state boundaries that are shared across participants. The other method is to average the data across participants before estimating state boundaries and the optimal number of states. Both of these methods assume that state boundaries are the same across participants. This assumption is true in the current simulation. We observed that both methods (averaging and crossvalidation) resulted in more accurate estimations of the number and the location of state boundaries. However, with high amounts of noise, leave-one-out cross validation still resulted in a strong overestimation of the number of state boundaries. Using 2-fold cross-validation or averaging half of the data resulted in similar levels of performance. Averaging the data across all 20 simulated participants showed the best performance, both for estimating the number of state boundaries and for identifying the location of state boundaries. When all data were averaged, the state boundaries could be detected accurately even when the noise SD was 10 times higher than the SD of the neural patterns that defined the states.

When data are averaged across multiple participants, or when using cross-validation, we assume that neural states are shared across participants. However, that assumption might not always be valid. In the simulation E, we examined whether it is possible to recover shared states when each subject also has some states that are not shared with the other participants. In particular, we simulated data in which participants always traversed 15 states. However, the proportion of states that was shared with other participants could vary. Specifically, we removed 10%, 20% or 40% of the group-level state boundaries in each participant and we randomly added the same number participant-specific states with unique activity patterns. In this simulation the amounts of noise in the data were kept low (SD=0.1)

We found that leave-one-out cross-validation performed poorly when the proportion of participant-specific states increased. In contrast, when we averaged the data across all participants, we were able to recover the simulated number of states correctly and we found a high similarity between the group-level simulated and estimated state boundaries (see figure 8).

**Figure 8.**
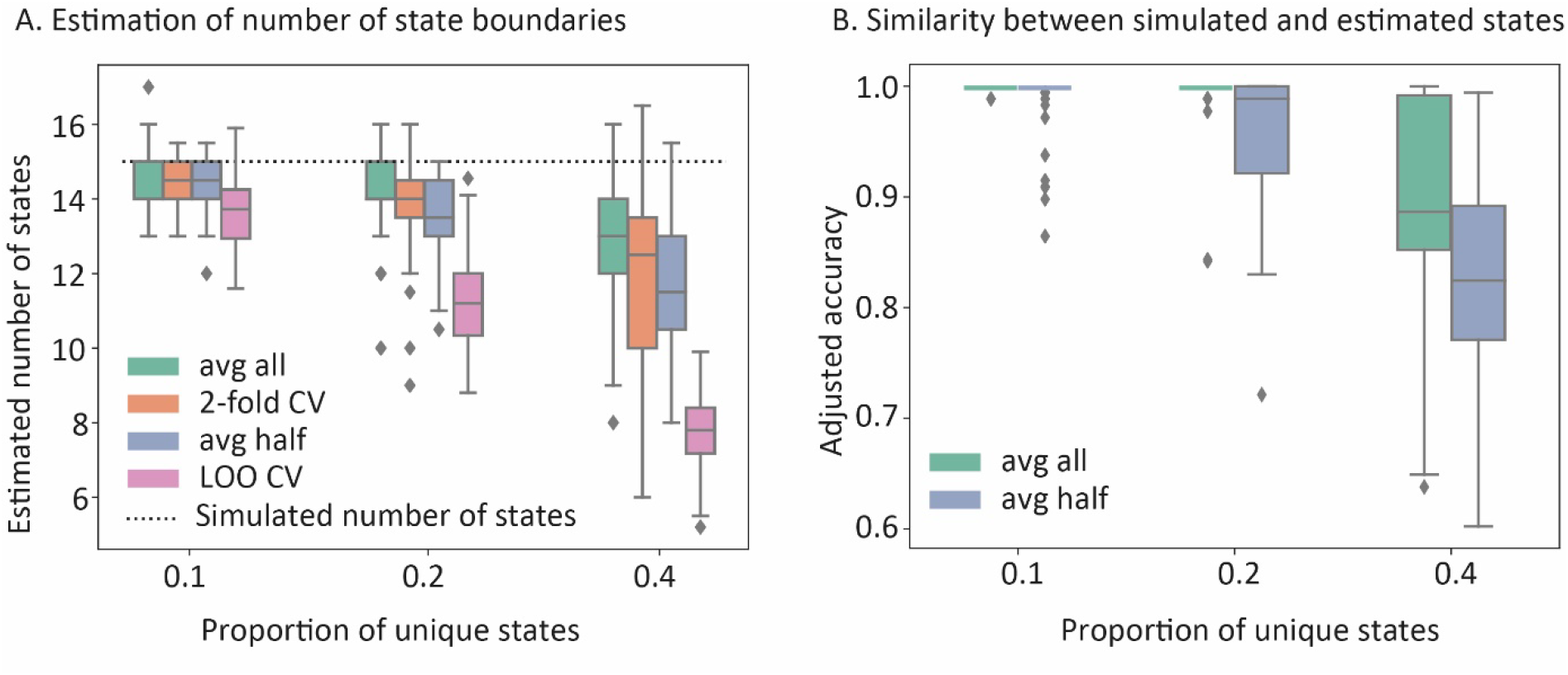
Results of simulation E. Comparing the effects of participant-specific states and data-averaging/cross validation on A) estimating the number of states B) the similarity between simulated and estimated state boundaries when we assume that the number of states (k=15) is known. avg all = averaging the data across all simulated participants; 2-fold CV = cross validation across 2 folds (independent groups). avg half = averaging the data in two independent groups; LOO CV = cross validation using the leave-one-out method.

Together, these simulations suggest that averaging the data across participants enables more reliable identification of neural state boundaries that are shared across the group, even if there is some degree of inter-individual variability in state boundary locations.

To investigate whether differences in the shape of the HRF might bias estimates of the number of neural states we performed an additional simulation (simulation F). In particular, we varied the HRF peak (4, 6 or 8 s) and the dispersion of the HRF (0.5, 1 or 2 s). We found that differences in the HRF shape did not affect the estimation of the number of states (see supplementary figure 4).

### Real data – data averaging

To get more insight into the optimal approach for analyzing real data, we compared different analysis choices in this final section. First, we investigated how data averaging affects the reliability of the recovered state boundaries and the estimated optimal number of states. Second, we investigated how these outcome measures are affected by the number of voxels in the searchlight.

In line with the results of our simulations, we found that the reliability of the state boundaries increased as the voxel timecourses were averaged over more participants (see figure 9A). For area IT and area V5, we found that the reliability increase tapered off when more than 25 participants were averaged. For AG, the mPFC and V1, we found that the reliability kept increasing as more participants were added. For single-participant data, the reliability was very low, suggesting that in this dataset, states cannot be estimated reliably in single participant data. We found that the estimate of the optimal number of states is reasonably stable across the number of averaged participants, as long as that number is around 17 or higher. Below that, we observe an increase in the estimated optimum (see figure 9B). This is similar to the results we observe in our simulations for data with low signal to noise, suggesting that the method starts to overestimate the number of state boundaries due to noisy data. This is particularly clear when data from single participants is used.

**Figure 9:**
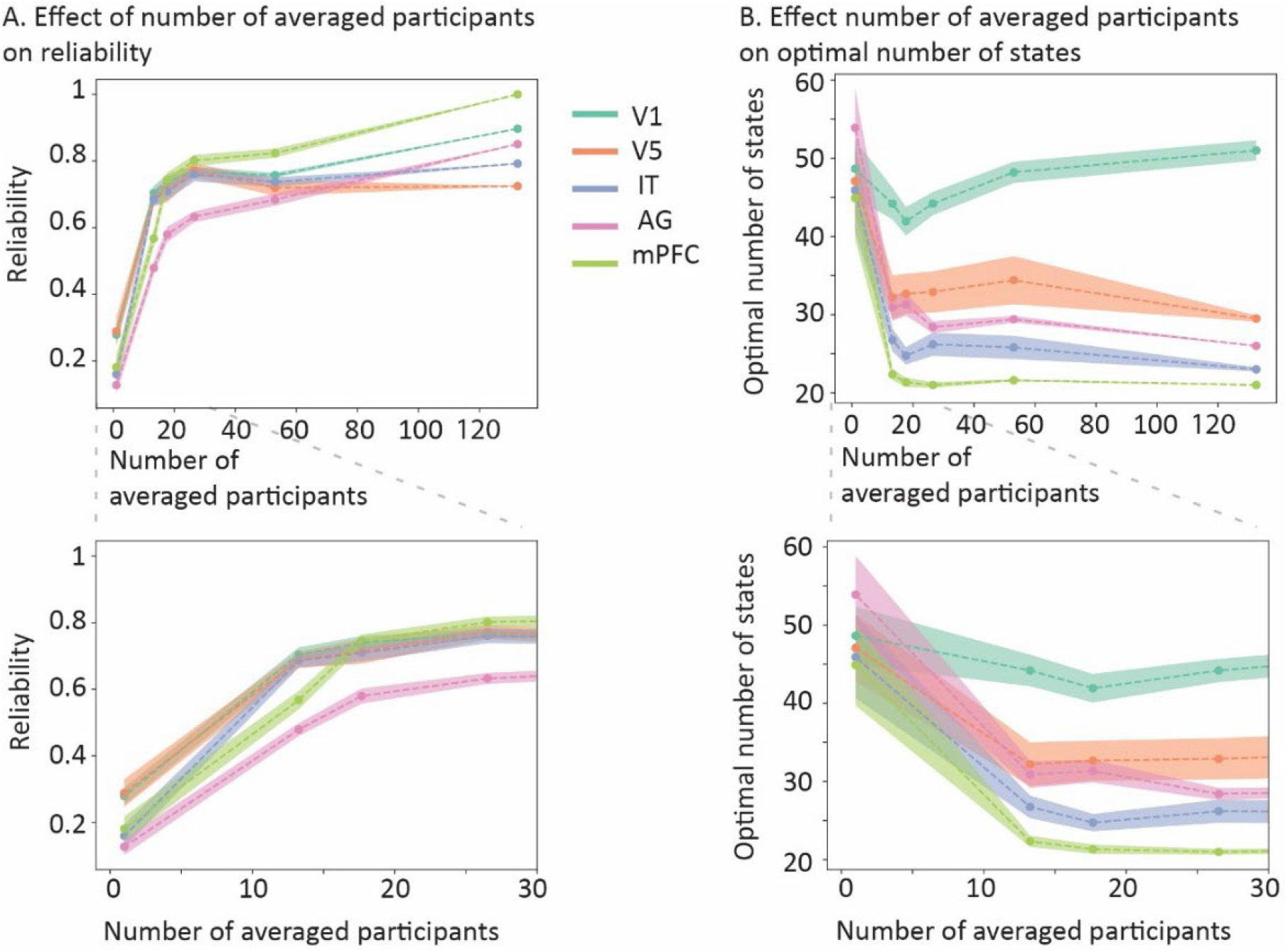
Investigating the effect of data averaging on A) the reliability of state boundaries and B) the estimated optimal number of states. The shaded area indicates the standard error across the independent (groups of) participants.

Averaging the voxel timecourses across participants allows us to isolate the BOLD signals that are evoked by the movie and shared across participants, from signals that have a non-neural origin (e.g. head motion and respiration) as well as neural signals that are not evoked by the movie. However, the disadvantage of this approach is that noise is introduced when voxels are not correctly aligned across participants. By using hyperalignment, we attempted to optimally align voxels. However, this alignment will never be perfect. Therefore, we also explored what happens when, instead of averaging voxel timecourses, we average the temporal correlation matrices, shown in figure 1B (panel 1). This would reduce effects of voxel misalignment, but also decrease the noise reduction effects that we get from voxel timecourse averaging. To compare the two methods, we investigated the reliability of the temporal correlation matrices across 15 independent groups of participants. Reliability was measured as the similarity of the flattened temporal correlation matrix observed in each group and a flattened correlation matrix averaged across the remaining groups of participants. We found that in each ROI, the reliability was higher when we averaged the voxel timecourses than when we averaged the temporal correlation matrices (see supplementary figure 5). These results support our interpretation that averaging voxel timecourses and focusing on the movie-evoked signals allows us to more accurately identify shared state boundaries across participants.

We also investigated how the reliability of neural states is affected by hyperalignment and removing voxels with low inter-subject synchrony (ISS). Previous research used a threshold of ISS > 0.25 to exclude voxels that were not synchronized across participants (Baldassano et al., 2017). Our results showed that hyperalignment drastically improves the reliability of the detected neural states, while thresholding voxels with low ISS had a smaller and more mixed effect, reliability decreasing for some brain regions and increasing for others (see supplementary figure 6).

Finally, we explored how different searchlight sizes affect the reliability and the optimal number of states (see supplementary figure 7). We found that the reliability is similar across different searchlight sizes. The estimate of the optimal number of states was stable for searchlights with a radius of 8 mm (+/- 80 voxels) or larger.

## Discussion

Event segmentation is an important mechanism that allows us to understand, remember and organize ongoing sensory input. Recent work has suggested that event segmentation can be linked to regional changes in neural activity patterns (neural state boundaries). Accurate methods for identifying neural state boundaries are important to allow further investigation of the neural basis of event segmentation and its link to the temporal processing hierarchy of the brain. In this paper, we have introduced simple and effective new methods for identifying the location of the boundaries between neural states as well as the number of neural states in a brain area; greedy state boundary search (GSBS) and t-distance.

We have used a comprehensive set of simulations as well as analyses of real fMRI data to show that these methods outperform an existing method based on an adapted version of HMMs (Baldassano et al., 2017). In addition, we have investigated the impact of noise on the estimation of the location and number of state boundaries, and how this can be mitigated by data averaging.

The GSBS method we introduce here, differs from the HMM-based state segmentation method in a number of important ways. First, the HMM-based method fits all the states in one go. This fitting procedure causes the method to tend to identify states with the same duration, as we observed in our simulations as well as our real data analyses. This problem is partly, but not completely resolved when the fit is optimized by splitting and merging states sequentially. In contrast, the GSBS method performs an exhaustive iterative search in which states are not biased toward a particular duration. Second, the HMM-based method identifies a new set of states for each value of *k* (the number of states), while the GSBS method performs an iterative search, such that all but one of the boundaries that are detected for *k* states are similar to the boundaries that are detected for *k+1* states (up to a fine-tuning step). Combined with the simplicity of the GSBS method, this results in an up to 170-fold increase in computations speed. This makes the GSBS method very suitable to detect state boundaries across many different brain regions in a whole-brain searchlight approach. The iterative nature of the GSBS method also results in an automatic ordering of state boundaries, as the strongest boundaries (the biggest change in neural patterns) will be detected first.

For both methods, a separate measure is needed to identify the optimal number of states. Here we introduced the t-distance metric, which maximizes a t-statistic that reflects the distance between the within-state correlations and the correlations between consecutive states. The t-distance was able to accurately recover the number of simulated states in the simulated data and also resulted in the expected temporal hierarchy for the empirical data (Hasson et al., 2015; Honey et al., 2012; Lerner et al., 2011). In contrast, the original WAC metric by Baldassano et al. (2017) tended to overestimate the number of states unless the number of states was much smaller than the number of timepoints. Most likely, this is due to the autocorrelation introduced by the HRF. Similarly, the log likelihood metric that is computed by the HMM, was unable to recover the number of states accurately in the simulations. Note that in the real data analyses that were presented by Baldassano et al. (2017), the number of states was constrained to be much smaller than the number of timepoints (max. 120 states in 50 minutes), which is perhaps the reason that they did not encounter the same problem with WAC.

One important benefit of the HMM-based method over the GSBS method is that the forward-backward algorithm can be used to identify states with similar activity patterns across different datasets, even when the durations of these states are not matched across datasets. This makes it possible to look at similarities across movies with comparable scenes (Baldassano et al., 2018) or between encoding and recall (Baldassano et al., 2017). To get the benefits of the accurate state boundary detection with GSBS in combination with the flexibility of the HMM-method, it is possible to estimate the initial set of neural state patterns using the GSBS method and to subsequently input these in the forward-backward algorithm to match state patterns across datasets.

GSBS was designed with the purpose to study the neural correlates of event segmentation, as event boundaries tend to co-occur with neural state boundaries in some parts of the cortical hierarchy (Baldassano et al., 2017). Indeed, we found that state boundaries in the mPFC were reliably associated with event boundaries. Because of this aim, GSBS is not optimized to identify recurrent states. Based on a wealth of previous literature on whole-brain functional connectivity states, it is likely that brain activity states within brain regions are also recurrent over time (Allen et al., 2014; Meer et al., 2020; Vidaurre et al., 2017). Researchers interested in this recurrence can use our method to segment the timeseries into an initial set of meaningful units. Subsequently clustering algorithms, such as Gaussian Mixture Models, can be used to identify states that reoccur over time.

In addition to introducing new methods, we also performed extensive simulations and empirical analyses to investigate the effect of noise on the state boundary estimations and to examine how averaging the data can mitigate this effect. Our simulations showed that high levels of noise (signal to noise ratio of 1/10) result in an overestimation of the number of states and poor reconstruction of the state boundary locations. When we assume that states are shared between participants, e.g. because they are watching the same movie, we find that averaging the data allows us to estimate the number and location of state boundaries correctly even in these very high noise regimes. Our empirical data analyses showed overestimation of the number of states when boundaries were estimated on non-averaged single subject data, but not when data were averaged over 17 or more participants. This suggests that the state-changes we identified were driven by the neural signal that was evoked by the movie rather than the ‘background’ neural signals that were not shared between participants. Another set of simulations showed that state changes can be detected reliably on the group-level even if there is some inter-individual variability in the states that are visited by participants.

The methods we introduced here were optimized specifically for estimating regional state boundaries in fMRI data. However, they are also applicable in other settings, such as investigating functional connectivity states (Allen et al., 2014; Damaraju et al., 2014; Wang et al., 2016) or investigating state boundaries in electrophysiological data (Borst and Anderson, 2015; Silva et al., 2019; Vidaurre et al., 2016).

To conclude, we have introduced a set of simple and computationally fast new methods that allow researchers to estimate state boundaries in fMRI data. These methods were validated using real and simulated data, giving us good insights in how they should be used to answer empirical questions. These methods will give researchers new, well-validated tools to investigate state-boundaries in neural data and to investigate the neural underpinnings of event segmentation.

## Acknowledgements

LG is supported by a Veni grant [451-16-013] from the Netherlands Organization for Scientific Research. MVG is supported by a Vidi grant [639.072.513] from the Netherlands Organization for Scientific Research. We thank Aya Ben-Yakov for providing data on the subjective event boundaries in the Cam-CAN movie dataset. Data collection and sharing for this project was provided by the Cambridge Centre for Ageing and Neuroscience (CamCAN). CamCAN funding was provided by the UK Biotechnology and Biological Sciences Research Council (grant number BB/H008217/1), together with support from the UK Medical Research Council and University of Cambridge, UK.

## Supplementary figures

**Supplementary figure 1:**
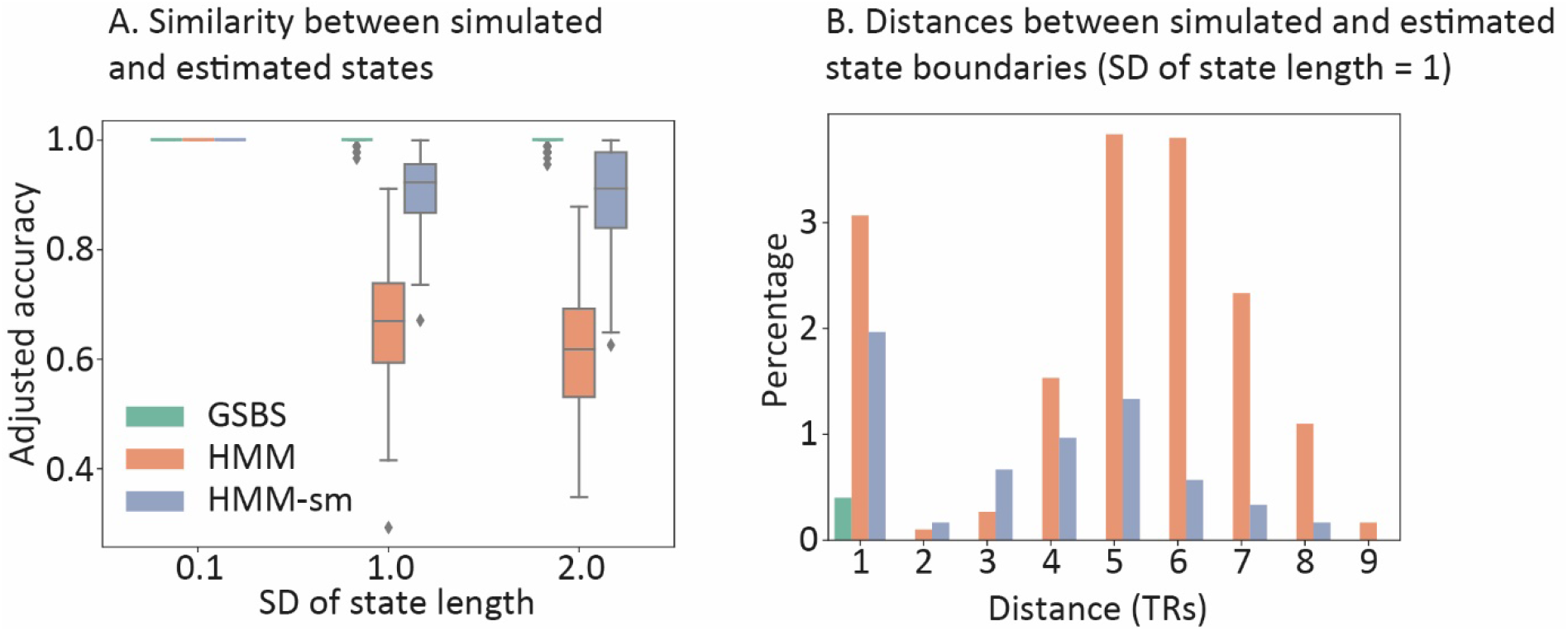
Results of simulation A. In this simulation the number of states is assumed to be known (k = 30). A) Similarity between simulated and estimated states for the HMM-based and GSBS methods. B) Distance between simulated and estimated state boundaries. The height of the bars indicates the percentage of the total number of estimated state boundaries for which the given distance was observed. Boundaries with a distance of zero (indicating perfect overlap between the simulated and estimates boundary) are not shown.

**Supplementary figure 2:**
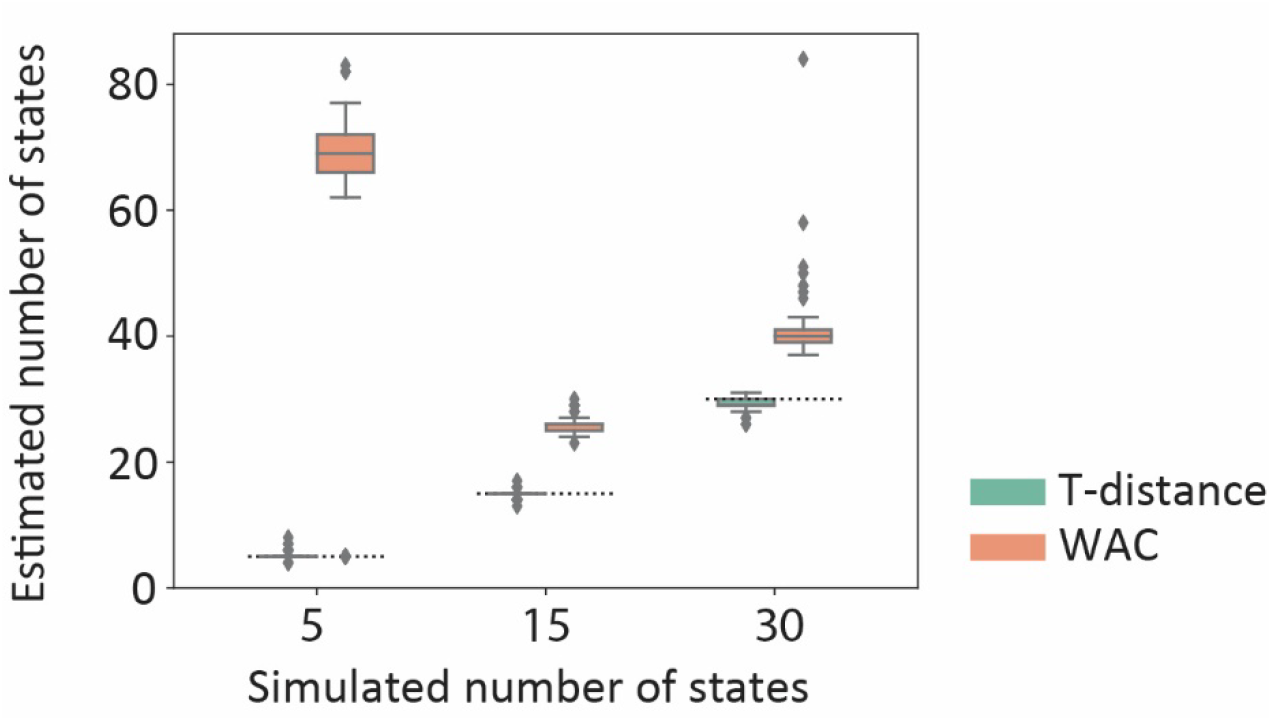
Results of simulation B. Comparison of the estimated and simulated number of states for T-distance and WAC. In this analysis, the 4 TRs around the diagonal of the correlation matrix were not taken into account in the computation of the WAC and t-distance metric (as in Baldassano et al. 2017). The dotted lines indicate the simulated number of states

**Supplementary figure 3:**
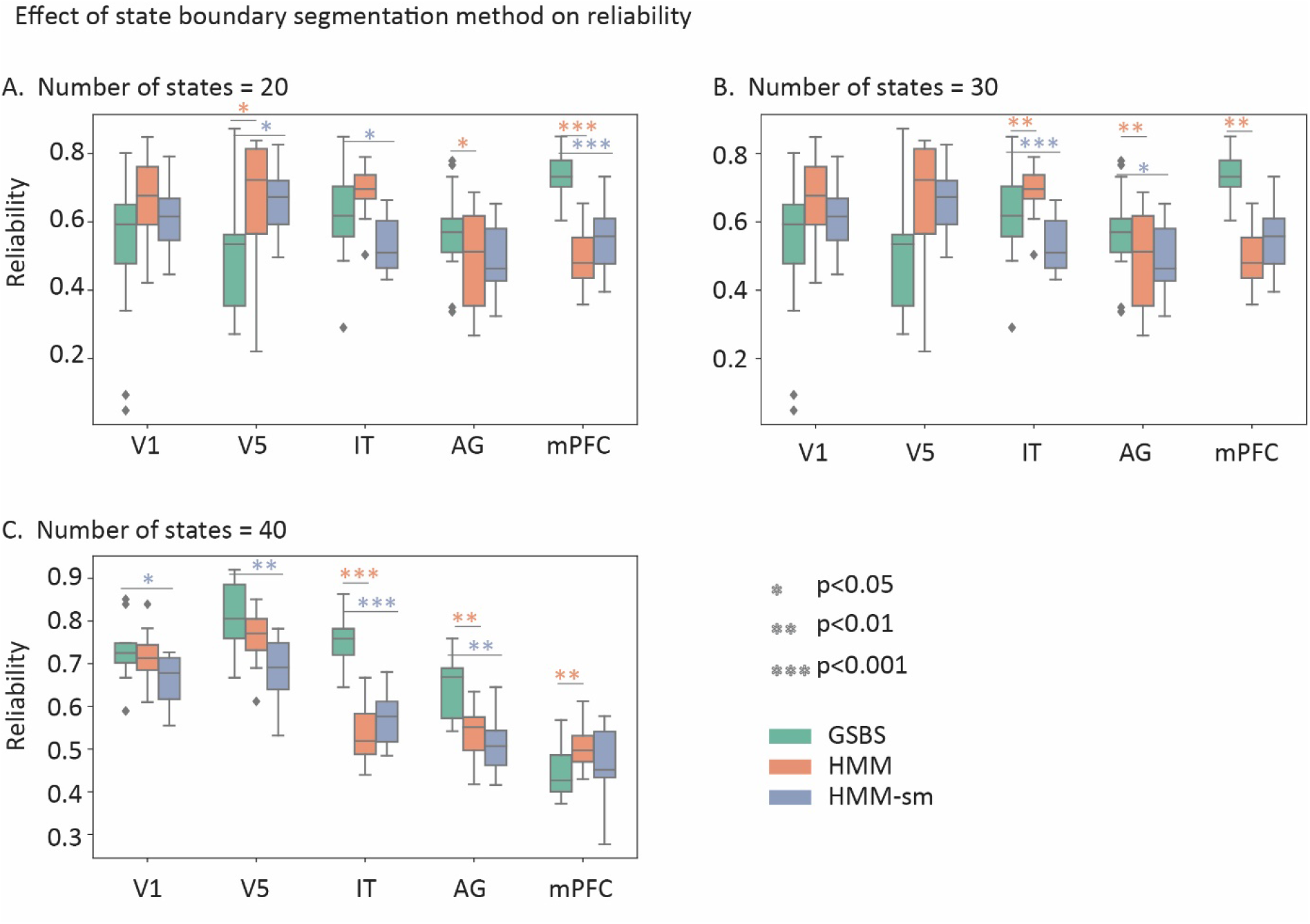
Reliability of state boundaries detected in real data using GSBS or HMM-based methods for different numbers of states.

**Supplementary figure 4.**
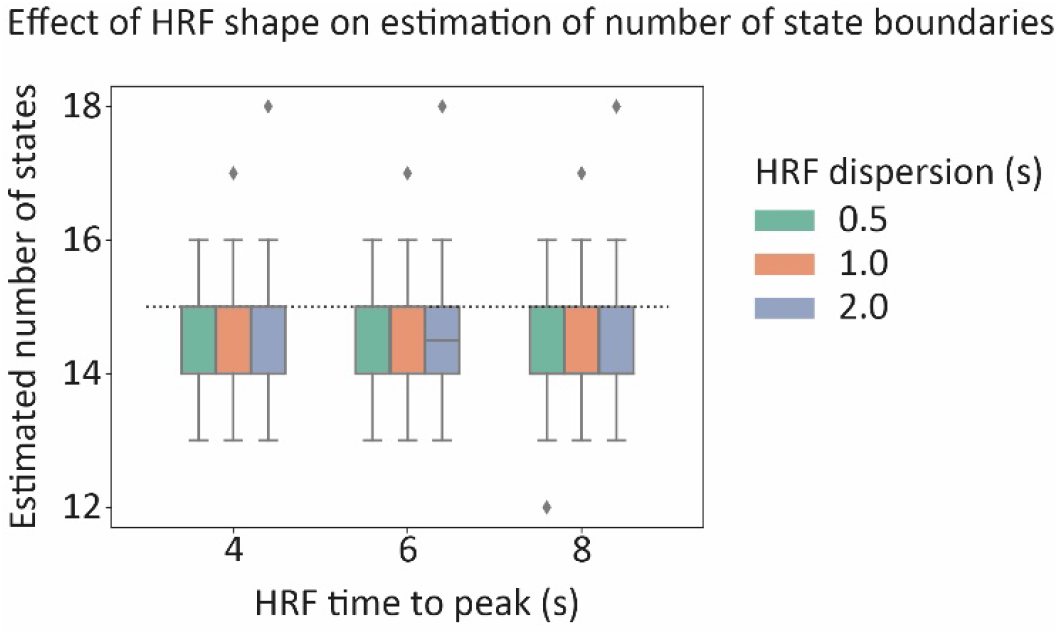
Results of simulation F. Effect of the HRF shape on the estimation of the number of state boundaries.

**Supplementary figure 5.**
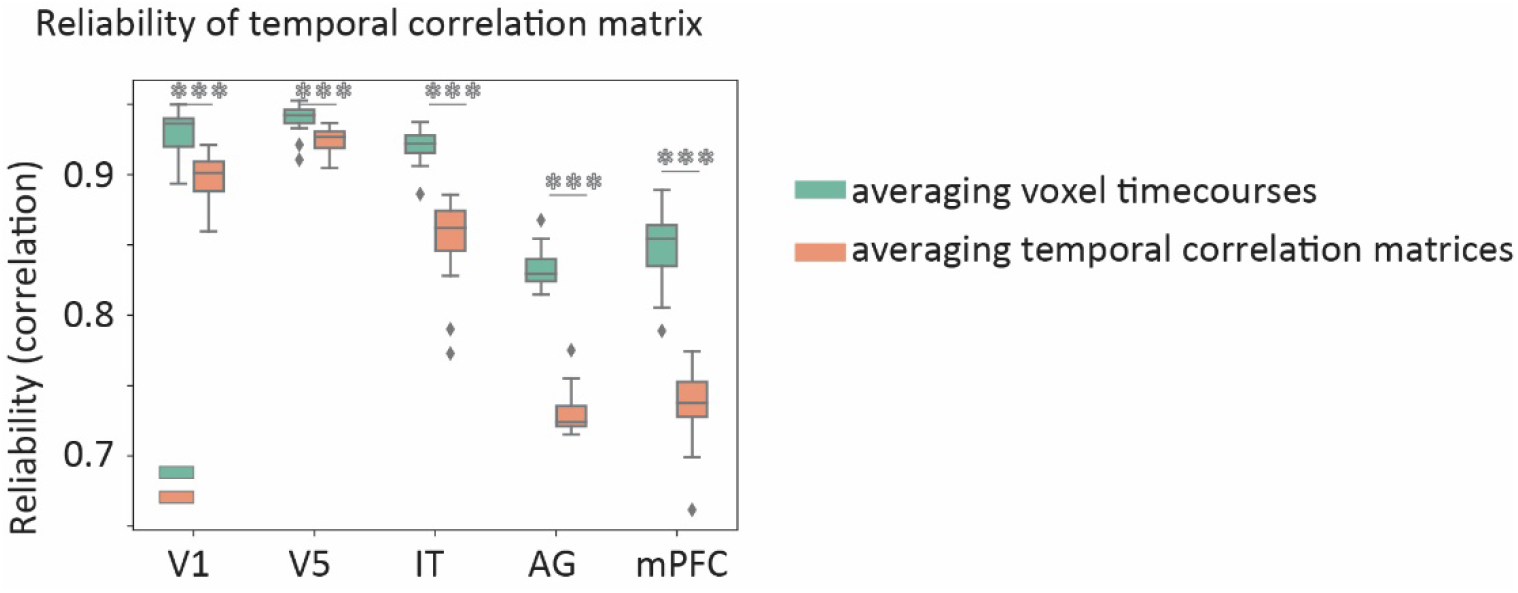
Effect of averaging voxel timecourses versus averaging temporal correlation matrices on the reliability of the temporal correlation matrix.

**Supplementary figure 6.**
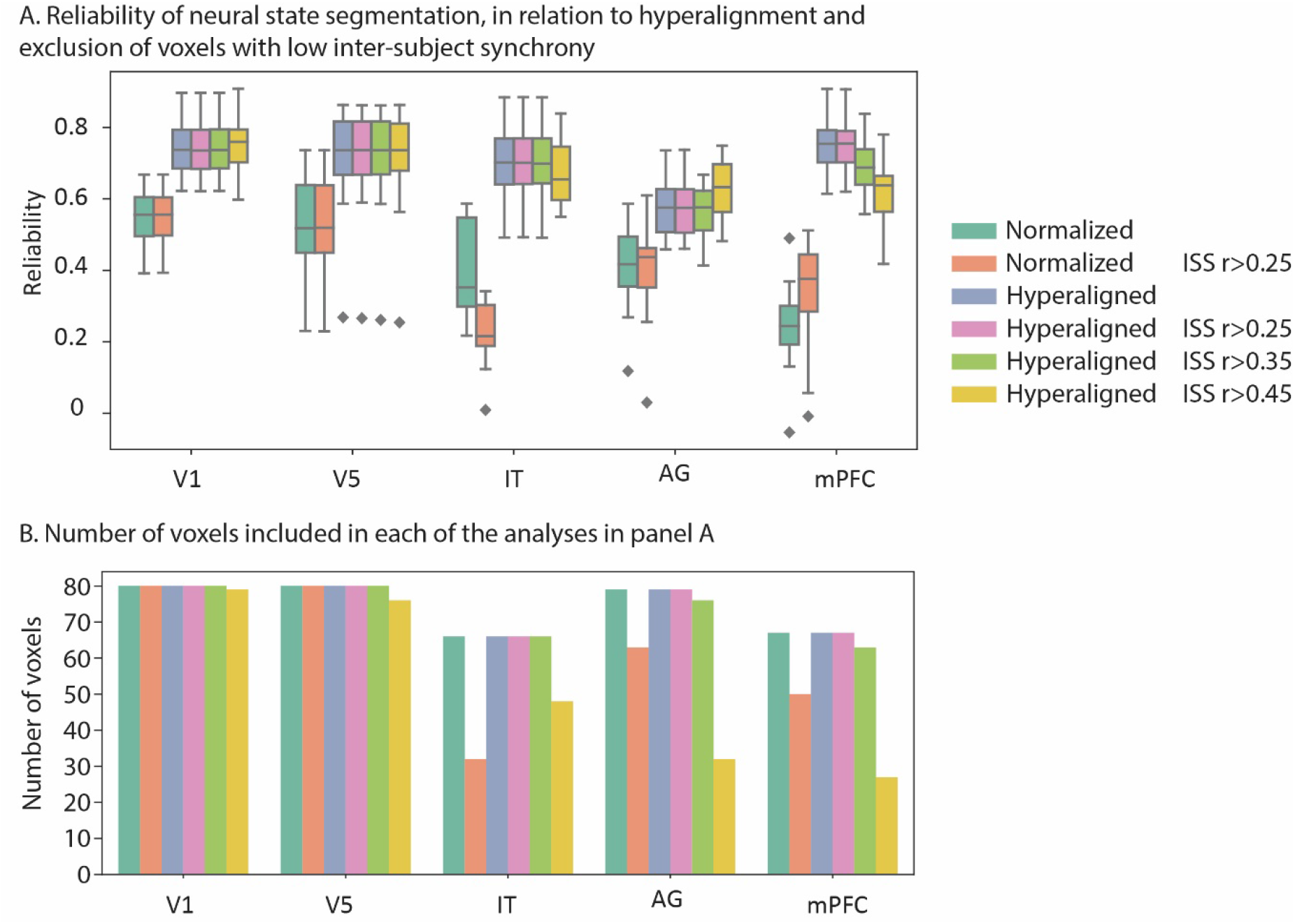
A) Effect of hyperalignment and voxel thresholding based on inter-subject synchrony (ISS) on the reliability of neural states. B) Number of voxels in each searchlight before and after thresholding.

**Supplementary figure 7.**
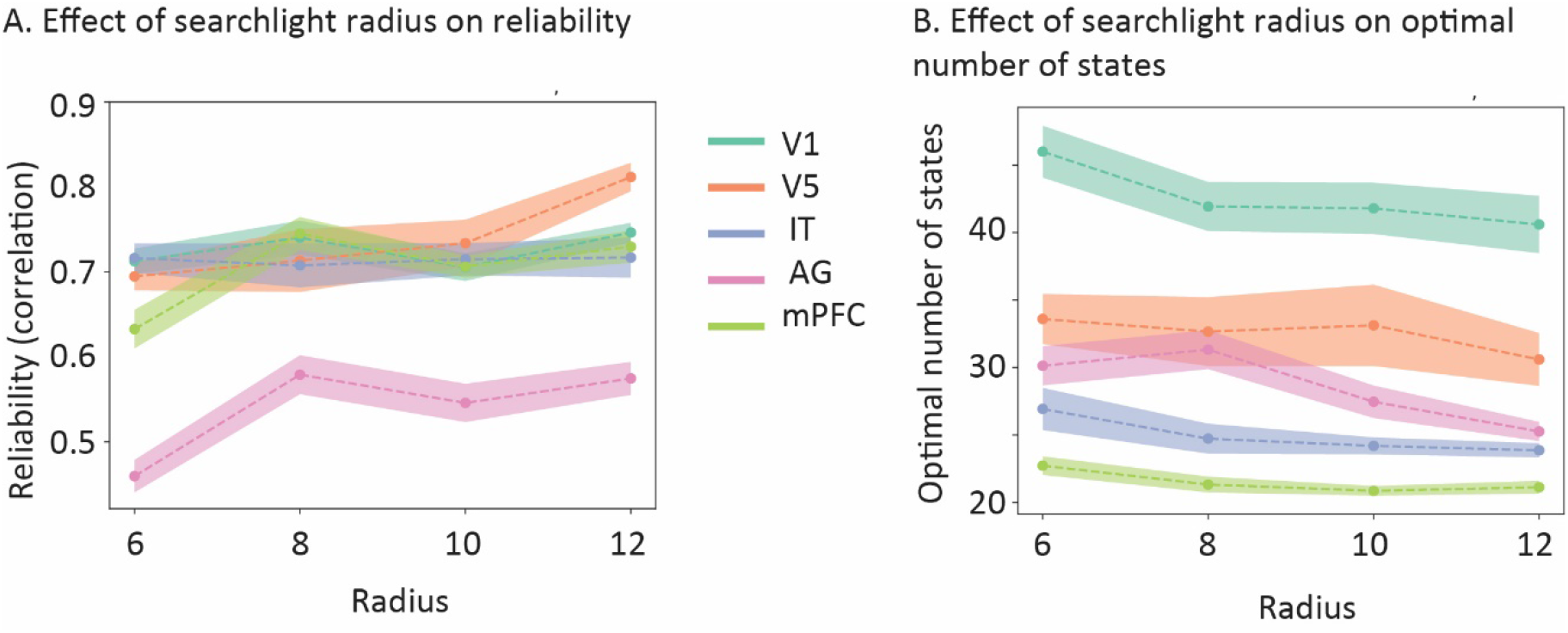
Effect of searchlight radius on A) reliability of state boundaries and B) optimal number of states. A radius of 6 mm corresponds to approximately 33 voxels, 8 mm is approximately 80 voxels, 10 mm is approximately 149 voxels and 12 mm is approximately 260 voxels.

## References

Allen, E.A., Damaraju, E., Plis, S.M., Erhardt, E.B., Eichele, T., Calhoun, V.D., 2014. Tracking whole-brain connectivity dynamics in the resting state. Cereb. Cortex 24, 663–76. https://doi.org/10.1093/cercor/bhs352

Ashburner, J., 2007. A fast diffeomorphic image registration algorithm. Neuroimage 38, 95–113.

Baldassano, C., 2020. Split-Merge HMMs [WWW Document]. URL http://www.chrisbaldassano.com/blog/2020/05/19/splitmerge/ (accessed 10.20.20).

Baldassano, C., Chen, J., Zadbood, A., Pillow, J.W., Hasson, U., Norman, K.A., 2017. Discovering Event Structure in Continuous Narrative Perception and Memory. Neuron 95, 709–721.e5. https://doi.org/10.1016/j.neuron.2017.06.041

Baldassano, C., Hasson, U., Norman, K.A., 2018. Representation of real-world event schemas during narrative perception. J. Neurosci. 38, 9689–9699. https://doi.org/10.1523/JNEUROSCI.0251-18.2018

Ben-Yakov, A., Henson, R.N., 2018. The Hippocampal Film Editor: Sensitivity and Specificity to Event Boundaries in Continuous Experience. J. Neurosci. 38, 10057–10068. https://doi.org/10.1523/JNEUROSCI.0524-18.2018

Borst, J.P., Anderson, J.R., 2015. The discovery of processing stages: Analyzing EEG data with hidden semi-Markov models. Neuroimage 108, 60–73. https://doi.org/10.1016/j.neuroimage.2014.12.029

Buonomano, D. V., Maass, W., 2009. State-dependent computations: spatiotemporal processing in cortical networks. Nat. Rev. Neurosci. 10, 113–125. https://doi.org/10.1038/nrn2558

Chang, L., Manning, J., Baldassano, C., Vega, A. de la Fleetwood, G., Geerligs, L., Haxby, J., Lahnakoski, J., Parkinson, C., Shappell, H., Shim, W.M., Wager, T., Yarkoni, T., Yeshurun, Y., Finn, E., 2020. naturalistic-data-analysis/naturalistic_data_analysis: Version 1.0. https://doi.org/10.5281/ZENODO.3937849

Chien, H.Y.S., Honey, C.J., 2020. Constructing and Forgetting Temporal Context in the Human Cerebral Cortex. Neuron 106, 675–686.e11. https://doi.org/10.1016/j.neuron.2020.02.013

Cox, R.W., 1996. AFNI: Software for analysis and visualization of functional magnetic resonance neuroimages. Comput. Biomed. Res. 29, 162–173. https://doi.org/10.1006/cbmr.1996.0014

Cribben, I., Haraldsdottir, R., Atlas, L.Y., Wager, T.D., Lindquist, M.A., 2012. Dynamic connectivity regression: determining state-related changes in brain connectivity. Neuroimage 61, 907–20. https://doi.org/10.1016/j.neuroimage.2012.03.070

Damaraju, E., Allen, E.A., Belger, A., Ford, J.M., McEwen, S., Mathalon, D.H., Mueller, B.A., Pearlson, G.D., Potkin, S.G., Preda, A., Turner, J.A., Vaidya, J.G., Van Erp, T.G., Calhoun, V.D., 2014. Dynamic functional connectivity analysis reveals transient states of dysconnectivity in schizophrenia. NeuroImage Clin. 5, 298–308. https://doi.org/10.1016/j.nicl.2014.07.003

DuBrow, S., Rouhani, N., Niv, Y., Norman, K.A., 2017. Does mental context drift or shift? Curr. Opin. Behav. Sci. 17, 141–146. https://doi.org/10.1016/j.cobeha.2017.08.003

Ezzyat, Y., Davachi, L., 2011. What constitutes an episode in episodic memory? Psychol. Sci. 22, 243–52. https://doi.org/10.1177/0956797610393742

Flores, S., Bailey, H.R., Eisenberg, M.L., Zacks, J.M., 2017. Event segmentation improves event memory up to one month later. J. Exp. Psychol. Learn. Mem. Cogn. 43, 1183–1202. https://doi.org/10.1037/xlm0000367

Geerligs, L., Cam-CAN Campbell, K.L., 2018. Age-related differences in information processing during movie watching. Neurobiol. Aging 72, 106:120. https://doi.org/10.1016/j.neurobiolaging.2018.07.025

Guntupalli, J.S., Hanke, M., Halchenko, Y.O., Connolly, A.C., Ramadge, P.J., Haxby, J. V, 2016. A Model of Representational Spaces in Human Cortex. Cereb. Cortex 26, 2919–2934. https://doi.org/10.1093/cercor/bhw068

Hanke, M., Halchenko, Y.O., Sederberg, P.B., Hanson, S.J., Haxby, J. V., Pollmann, S., 2009. PyMVPA: a Python Toolbox for Multivariate Pattern Analysis of fMRI Data. Neuroinformatics 7, 37–53. https://doi.org/10.1007/s12021-008-9041-y

Hasson, U., Chen, J., Honey, C.J., 2015. Hierarchical process memory: Memory as an integral component of information processing. Trends Cogn. Sci. 19, 304–313. https://doi.org/10.1016/j.tics.2015.04.006

Hasson, U., Malach, R., Heeger, D.J., 2009. Reliability of cortical activity during natural stimulation. Trends Cogn. Sci. 14, 40–48. https://doi.org/10.1016/j.tics.2009.10.011

Honey, C.J., Thesen, T., Donner, T.H., Silbert, L.J., Carlson, C.E., Devinsky, O., Doyle, W.K., Rubin, N., Heeger, D.J., Hasson, U., 2012. Slow cortical dynamics and the accumulation of information over long timescales. Neuron 76, 423–34. https://doi.org/10.1016/j.neuron.2012.08.011

Kiebel, S.J., Daunizeau, J., Friston, K.J., 2008. A Hierarchy of Time-Scales and the Brain. PLoS Comput. Biol. 4, e1000209. https://doi.org/10.1371/journal.pcbi.1000209

Kuhn, H.W., 1955. The Hungarian method for the assignment problem. Nav. Res. Logist. Q. 2, 83–97. https://doi.org/10.1002/nav.3800020109

Kundu, P., Inati, S.J., Evans, J.W., Luh, W.-M., Bandettini, P.A., 2012. Differentiating BOLD and non-BOLD signals in fMRI time series using multi-echo EPI. Neuroimage 60, 1759–70. https://doi.org/10.1016/j.neuroimage.2011.12.028

Kundu, S., Ming, J., Pierce, J., McDowell, J., Guo, Y., 2018. Estimating dynamic brain functional networks using multi-subject fMRI data. Neuroimage 183, 635–649. https://doi.org/10.1016/j.neuroimage.2018.07.045

Kurby, C.A., Zacks, J.M., 2008. Segmentation in the perception and memory of events. Trends Cogn. Sci. 12, 72–79. https://doi.org/10.1016/j.tics.2007.11.004

Lerner, Y., Honey, C.J., Silbert, L.J., Hasson, U., 2011. Topographic Mapping of a Hierarchy of Temporal Receptive Windows Using a Narrated Story. J. Neurosci. 31, 2906–2915. https://doi.org/10.1523/JNEUROSCI.3684-10.2011

Meer, J.N. van der Breakspear, M., Chang, L.J., Sonkusare, S., Cocchi, L., 2020. Movie viewing elicits rich and reliable brain state dynamics. Nat. Commun. 11, 1–14. https://doi.org/10.1038/s41467-020-18717-w

Newtson, D., Engquist, G.A., Bois, J., 1977. The objective basis of behavior units. J. Pers. Soc. Psychol. 35, 847–862. https://doi.org/10.1037/0022-3514.35.12.847

Sargent, J.Q., Zacks, J.M., Hambrick, D.Z., Zacks, R.T., Kurby, C.A., Bailey, H.R., Eisenberg, M.L., Beck, T.M., 2013. Event segmentation ability uniquely predicts event memory. Cognition 129, 241–255. https://doi.org/10.1016/j.cognition.2013.07.002

Shafto, M.A., Tyler, L.K., Dixon, M., Taylor, J.R., Rowe, J.B., Cusack, R., Calder, A.J., Marslen-Wilson, W.D., Duncan, J., Dalgleish, T., Henson, R.N., Brayne, C., Cam-CAN Matthews, F.E., 2014. The Cambridge Centre for Ageing and Neuroscience (Cam-CAN) study protocol: A cross-sectional, lifespan, multidisciplinary examination of healthy cognitive ageing. BMC Neurol. 14. https://doi.org/10.1186/s12883-014-0204-1

Silva, M., Baldassano, C., Fuentemilla, L., 2019. Rapid Memory Reactivation at Movie Event Boundaries Promotes Episodic Encoding. J. Neurosci. 39, 8538–8548. https://doi.org/10.1523/JNEUROSCI.0360-19.2019

Taylor, J.R., Williams, N., Cusack, R., Auer, T., Shafto, M. a., Dixon, M., Tyler, L.K., Cam-CAN Henson, R.N., 2017. The Cambridge Centre for Ageing and Neuroscience (Cam-CAN) data repository: Structural and functional MRI, MEG, and cognitive data from a cross-sectional adult lifespan sample. Neuroimage 144, 262–269. https://doi.org/10.1016/j.neuroimage.2015.09.018

Truong, C., Oudre, L., Vayatis, N., 2020. Selective review of offline change point detection methods. Signal Processing 167, 107299. https://doi.org/10.1016/j.sigpro.2019.107299

Vidaurre, D., Quinn, A.J., Baker, A.P., Dupret, D., Tejero-Cantero, A., Woolrich, M.W., 2016. Spectrally resolved fast transient brain states in electrophysiological data. Neuroimage 126, 81–95. https://doi.org/10.1016/j.neuroimage.2015.11.047

Vidaurre, D., Smith, S.M., Woolrich, M.W., 2017. Brain network dynamics are hierarchically organized in time. Proc. Natl. Acad. Sci. 114, 12827–12832. https://doi.org/10.1073/pnas.1705120114

Wang, C., Ong, J.L., Patanaik, A., Zhou, J., Chee, M.W.L., 2016. Spontaneous eyelid closures link vigilance fluctuation with fMRI dynamic connectivity states. Proc. Natl. Acad. Sci. U. S. A. 113, 9653–9658. https://doi.org/10.1073/pnas.1523980113

Xu, Y., Lindquist, M.A., 2015. Dynamic connectivity detection: an algorithm for determining functional connectivity change points in fMRI data. Front. Neurosci. 9. https://doi.org/10.3389/fnins.2015.00285

Yarkoni, T., Poldrack, R.A., Nichols, T.E., Van Essen, D.C., Wager, T.D., 2011. Large-scale automated synthesis of human functional neuroimaging data. Nat. Methods 8, 665–670. https://doi.org/10.1038/nmeth.1635

Zacks, J.M., Speer, N.K., Swallow, K.M., Braver, T.S., Reynolds, J.R., 2007. Event perception: A mind-brain perspective. Psychol. Bull. 133, 273–293. https://doi.org/10.1037/0033-2909.133.2.273

Zacks, J.M., Speer, N.K., Vettel, J.M., Jacoby, L.L., 2006. Event understanding and memory in healthy aging and dementia of the Alzheimer type. Psychol. Aging 21, 466–482. https://doi.org/10.1037/0882-7974.21.3.466

Zacks, J.M., Tversky, B., Iyer, G., 2001. Perceiving, remembering, and communicating structure in events. J. Exp. Psychol. Gen. 130, 29–58.

